# *De novo* assembly of the black flounder genome. Why do pleuronectiformes have such a small genome size?

**DOI:** 10.1101/2023.03.27.534153

**Authors:** Fernando Villarreal, Germán F. Burguener, Ezequiel J. Sosa, Nicolas Stocchi, Gustavo M. Somoza, Adrián Turjanski, Andrés Blanco, Jordi Viñas, Alejandro S. Mechaly

## Abstract

Black flounder (*Paralichthys orbignyanus*) is an economically important marine fish with aquaculture potential in Argentina due to its market value. In this study, we sequenced the whole genome using an Illumina sequencing technology. We started with two independent libraries (from one female and one pool of females; each with 150 bp paired-end reads, a mean insert length of 350 bp, and >35 X-fold coverage). Each library was assembled separately using SOAPdenovo2 and the resulting contigs were scaffolded with SSPACE3 before gaps were filled with GapCloser. In vertebrates, including teleosts, the number of transposable elements (TEs) is related to genome size, but it remains unclear whether the size of introns and exons also plays a role. Therefore, the main objective of the present study was to test whether the small genome size of Pleuronectiformes is related to the size of their introns and exons. The assemblies resulted in a genome size of ∼538 Mbp (41.35% GC content, 0.11% undetermined bases). Analysis of the assemblies at the core genes level (subset of the 458 universally expressed KOG families) revealed that more than 98% of core genes are present, with more than 78% of them having more than 50% coverage. This indicates a fairly complete and accurate genome at the coding sequence level. Prediction of genes based on statistical predictors (geneid) and sequence-based predictors (Exonerate, using a closely related species, *Paralichthys olivaceus*, as a reference) was performed. This revealed 25,231 protein-coding genes, 445 tRNAs, 3 rRNAs, and more than 1,500 non-coding RNAs of other types (including a complete set of spliceosomes and several types of snoRNA and miRNA). As a result, this study concluded that the reduced genome size of flounders is related to a reduction in transcript size, mainly through a reduction in exon number, but also through a reduction in large introns. Thus, both components seem to be involved in the strategy of genome reduction in Pleuronectiformes.

## INTRODUCTION

Pleuronectiforms are an interesting order of fish due to their adaptations to demersal life and the presence of a drastic metamorphosis from bilateral pelagic larval symmetry to flatfish symmetry in the adult stage (Robledo et al., 2017; Lü et al., 2021). This order consists of a total of 772 species distributed among 129 genera and 14 families (Nelson et al., 2016). Some of the species are of great economic importance due to their importance to fisheries and aquaculture (Seitz et al., 2008). Within this order, the genus *Paralichthys* is one of the most relevant species for aquaculture. In Asia, aquaculture of the Japanese flounder (*Paralichthys olivaceus*) is well established (Seikai, 2002). In Latin America, other species belonging to this genus have also been identified with potential for aquaculture. Worth mentioning because of their importance to fisheries and aquaculture are Chilean flounder (*Paralichthys adspersus*), endemic to Peru and Chile, and black flounder (*Paralichthys orbignyanus*) in Brazil, Argentina, and Uruguay (Díaz de Astarloa, 2002).

Black flounder inhabit shallow estuaries and coastal waters from Rio de Janeiro (Díaz de Astarloa, 2002) to the Gulf of San Matías in northern Patagonia (Díaz de Astarloa and Munroe, 1998). As this species is a promising candidate for aquaculture, numerous zootechnical studies have been conducted on it in the last two decades (Bambill et al., 2006; Boccanfuso et al., 2019; López et al., 2009; Magnone et al., 2015; Sampaio and Bianchini, 2002; Sampaio et al., 2007; Radonic et al., 2007; Radonic and Macchi, 2009). However, there is an important knowledge gap in the genetic resources for this species. For example, only 12 protein sequences are deposited in the GenBank® sequence database (http://www.ncbi.nlm.nih.gov/). This is even more remarkable considering that genetics and genomics are considered one of the most important pieces of information for the development of the aquaculture and fisheries industry (Cerdà et al., 2010). This is particularly important for the study of sex determination and differentiation, sex ratio, nutrition, reproduction, genetic population structure, and other important biological traits. In this context, a reference genome is an essential requirement for the study of all these processes and mechanisms.

In the last decade, next-generation sequencing (NGS) technology has gained acceptance and played a critical role in obtaining whole genome sequences from different of non-model fish species. According to the *Ensembl* Genome Browser (https://www.ensembl.org), hundreds of fish species with whole genome sequences (WGS) are available to date, and the number of species whose genomes have been sequenced continuous increasing. Among Pleuronectiformes, for example, thirteen species have their genome sequenced: tongue sole (*Cynoglossus semilaevis*) (Chen et al., 2014), Turbot (*Scophthalmus maximus*) (Maroso et al., 2018), Senegalese sole (*Solea senegalensis*) (Guerrero-Cózar, et al., 2021), Japanese flounder (*Paralichthys olivaceus*) (Shao et al., 2017), Spotted halibut (*Verasper variegatus*) (Zhao et al., 2021), Hogchoker (*Trinectes maculatus)*, Pelican flounder (*Chascanopsetta lugubris)*, Oriental sole (*Brachirus orientalis)*, Bloch’s tonguesole (*Paraplagusia blochii)*, New Zealand turbot (*Colistium nudipinnis)*, Ocellated flounder (*Pseudorhombus dupliocellatus)*, Starry flounder (*Platichthys stellatus)*, and Indian halibut *Psettodes erumei*) (Lü et al., 2021). In addition, recent studies have compiled the genomes at the chromosome level of turbot (Martínez et al., 2021) and spotted halibut (Zhao et al., 2021). A recent study also analyzed the origin of the specialized body structure of flatfishes, the genomes of 11 flatfish species representing 9 of the 14 pleuronectiform families of this order were sequenced (Lü et al., 2021). In this frame, sequencing and annotation of the black flounder genome would expand the knowledge of this species so that alternative genomic methods could be used to improve fish breeding and to conduct comparative and evolutionary studies with other flatfish species.

Comparative genomics has revealed striking differences in genome size even in closely related species. In teleosts, this variation is exceptional, with enormous variance not only in genome size but also in chromosome number (Mank and Avise, 2006; Sarropoulou and Fernandes, 2011; Yuan et al., 2018). Previous studies have clearly shown that the Pleuronectiformes, along with the pufferfishes, seahorses, pipefishes, and Anabantiformes have less DNA compared to other teleost groups (Hinegardner, 1968; Hinegardner and Rosen, 1972), suggesting that they have smaller genomes. One of the hypotheses linked a small genome is the reduction in size of exons, and especially introns (Zhang and Edwards, 2012). Alternatively, the frequency of repetitive elements was positively correlated with genome size (Yuan et al., 2018). Pleuronectiform flatfish have only less than 9.0% of repetitive elements (Aparicio et al., 2002), compared to up to 60% in the salmon genome (Lien et al., 2016). However, the causes and consequences of this putative reduction in the genomes of these fish are still far from clear. In general, the variations in genome size are explained by the balance between genome-level mechanisms that lead to genome enlargement (duplication, transposable elements, and polyploidy) and genetic mechanisms that lead to genome reduction (deletions and DNA repair mechanisms) (Brainerd et al., 2001).

Thus, the main goal of the present study was to perform the first sequencing and characterization of the whole genome of black flounder. And therefore, to proportionate an important genomic tool for further research on this species. In addition, comparative genome analysis was used to determine whether Pleuronectiformes, present a small genome size compared to other teleost species. The analysis compares the exon and intron size of genes in different Pleuronectiformes in relation with homologous genes in the other teleost genomes and also considers the number of repetitive elements.

## MATERIAL AND METHODS

### Fish sampling and DNA extraction

Adult black flounder, *Paralichthys orbignyanus* were obtained from the Estación Experimental de Maricultura (INIDEP, Argentina). Fin tissue samples were collected and preserved in 96% ethanol. Genomic DNA was extracted from one adult female and a pool of females (n=10). Samples were lysed in 300 μl SSTNE extraction buffer (Blanquer, 1990) with SDS (0.1%) and 5 μl proteinase K (20 mg/ml) for 3 h at 55 °C. After 20 min at 70 °C, samples were treated for RNA digestion with 7.5 μl RNAse (10 mg/ml) for 1 h at 37 °C. Total DNA was purified with cold absolute ethanol (1 ml) after protein precipitation with 5 M NaCl. DNA quality (high molecular weight > 20 kb) was first assessed on agarose gels and DNA quantity was measured using NanoDrop® ND-1000 spectrophotometer (NanoDrop® Technologies Inc). Finally, DNA concentration was accurately measured using a Qubit fluorometer (Life Technologies). Fish were handled in accordance with internal institutional regulations (CICUAE-INIDEP).

### Genome sequencing and genome assemblies

All samples were adjusted to 300 ng/uL and fragmented by ultrasonication on Covaris, aiming for a fragment size of 300 bp. Barcoded libraries were prepared using Wafergen’s PrepX ILM DNA Library Kit on Apollo and beads size selected before being pooled and sequenced at 150 bp at the paired end on HiSeq-4000. All procedures followed the manufacturer’s instructions. We performed multiple QC sets during library preparation using 300 ng of human gDNA samples as internal control and all QC passed for this project.

Two Illumina paired-end libraries were constructed (from one female and one pool of females) and sequenced with a read length of 150 bp and an average fragment size of ∼350 bp. Reads were first filtered using Trimmomatic (Bolger et al., 2014) and PrinSeq (Bolger et al., 2014; Schmieder and Edwards, 2011) to remove adapters and low-quality reads, and then sequencing errors were corrected using Quake (Kelley et al., 2010).

The libraries were assembled individually using SOAPdenovo 2.04 (Luo et al., 2012) with a kmer size of 29, gap filled with GapCloser (Boetzer and Pirovano, 2012) and finally scaffolded with SSPACE3 (Boetzer et al., 2011) with default parameters.

### Genome annotation

First, genomes were masked with Repeatmasker (Hancock, 2004) using the Repbase (Bao et al., 2009) and DFam (Hubley et al., 2016) databases. Using the masked scaffolds, we searched for t-RNAs, r-RNAs and nc-RNAs using tRNAScan-SE (Zou et al., 2015), RNAmmer (Lagesen et al., 2007), and Infernal (Nawrocki and Eddy, 2013) + RFAM (Griffiths-Jones 2003), respectively. Then, we performed gene prediction using GeneId (Blanco and Abril 2009) and Exonerate (Slater and Birney, 2005) trained with Uniprot/Swissprot (Pundir et al., 2016) and the Japanese flounder (*Paralichthys olivaceus*) genome (Shao et al., 2017). All proteins were annotated using InterProScan (Hancock and Bishop, 2004).

### Gene family clustering and validation

To find orthologous proteins in other fish species, we downloaded the genome data of *D. rerio* (Howe et al., 2013), *C. semilaevis* (Chen et al., 2014), *S. maximus* (Xu et al., 2020), *P. olivaceus* (Shao et al., 2017 from the Zebrafish Genome Project and *Ensembl*. All protein sequences were aligned using BLASTP (Altschul et al., 1990) with an E value cutoff less than 1e−5. BLASTP results were parsed and imported into a MySQL database. Tables within were created by OrthoMCL (Li et al., 2003) to identify homologues with thresholds of percentMatchCutoff = 50 and evalueExpo-nentCutoff = 1e−6. To evaluate our annotation, we used Benchmarking Universal Single-Copy Orthologs (BUSCO, v3.0.2)37 to assess the assembled genome sequences. We used BUSCO with single copy orthologues from actinopterygii_odb10 to assess the completeness of the genome assembly and annotation.

### Identification and comparison of Transposable elements

Transposable elements (TEs) were quantified for each genome applying a homology-based approach using RepeatMasker (Tarailo-Graovac et al., 2009) with Actinopterygii species repeats and default parameters. Relationship between genome size and TEs percentage in the genome were plotted using ggplot2 R package, and correlations were calculated by Spearman method.

### Analysis of C-value in teleosts

All fish C-values (in pg) available were retrieved from the Animal Genome Size Database (http://www.genomesize.com) (Gregory, 2020). As of March 2023, the website contained data for more than 1800 fish species from more than 70 orders (48 entries correspond to Pleuronectiformes species). When necessary, the average C-value per species was calculated. The C-value for *P. orbignyanus* was estimated assuming that 1 pg = 978 Mb (Doležel et al., 2003). Boxplots were generated using Plotly (Plotly Technologies Inc. Collaborative data science. Montréal, QC). Because most data sets were not normally distributed (according to the Shapiro-Wilk test), C-value data were analyzed with the Mann-Whitney test, comparing Pleuronectiformes with each order using JASP (version 0.16, JASP Team, 2021).

### Gene features size analysis at whole genome scale in teleost orders

Complete genome annotations were retrieved from ENSEMBL in gff3 format. We selected species in orders phylogenetically close to Pleuronectiformes. A total of twenty seven species representatives from ten fish orders were used: Anabantiformes (*Anabas testudineus*, *Betta splendens*, *Mastacembelus armatus*), Carangiformes (*Echeneis naucrates*, *Seriola dumerili*, *Seriola lalandi dorsalis*), Cichliformes (*Astatotilapia calliptera*, *Oreochromis niloticus*, *Pundamilia nyererei*), Cypriniformes (*Cyprinus carpio*, *Danio rerio*, *Sinocyclocheilus rhinocerous*), Cyprinodontiformes (*Fundulus heteroclitus*, *Kryptolebias marmoratus*, *Poecilia formosa*), Perciformes (*Cyclopterus lumpus*, *Gasterosteus aculeatus*, *Sander lucioperca*), Pleuronectiformes (*Cynoglossus semilaevis*, *Paralichthys orbignyanus* female, *Scophthalmus maximus*), Salmoniformes (*Hucho hucho*, *Oncorhynchus mykiss*, *Salmo trutta*), Labriformes (*Labrus bergylta*) and Tetraodontiformes (*Takifugu rubripes*, *Tetraodon nigroviridis*). See **Supplementary Table Ss1** for more details. Using a tailored python script, we retrieved the size of each gene (genomic locus) by calculating the difference between the final and initial base coordinates. For each transcript model at each locus, we calculated the number of exons and the size of each exon. As well as the transcript size by calculating the sum of the exon sizes. Finally, intron sizes were inferred by calculating the difference between the first base coordinate of an exon and the last base coordinate of the preceding exon.

The size distribution of exons and introns at the genome level were plotted using the kernel density estimation of the Seaborn package (parameters: gaussian kernel, band width 0.1 for exons and 0.01 for introns, and grid size of 1000 for exons and 5000 for introns). Boxplots for the size distributions were plotted using Plotly.

## Results

### Genome annotation

After filtering, the remaining reads accounted for a coverage of more than 35 X-fold on each genome and were retained for assembly (**Table 1**).

**Table 1:**
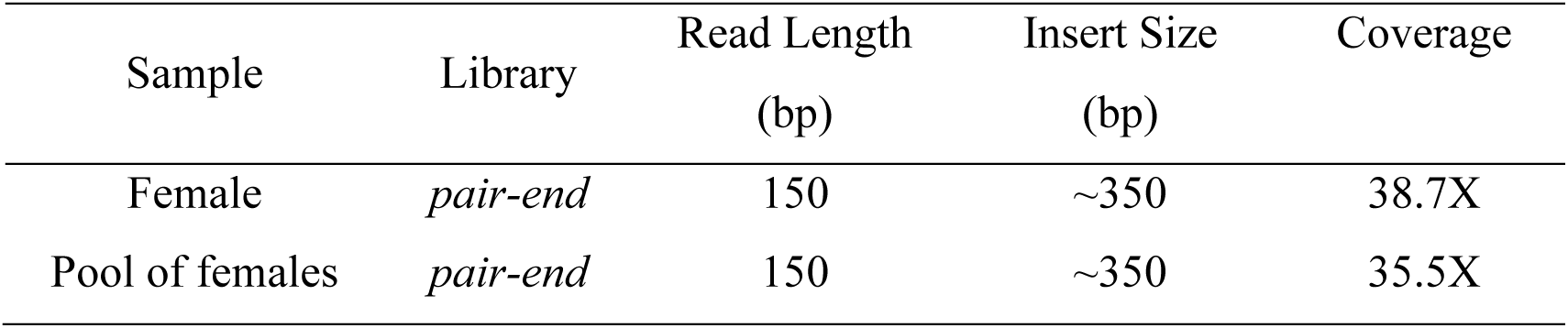
Genome coverage for each library. Genome sequencing (post-filtering)

| Sample | Library | Read Length<br>(bp) | Insert Size<br>(bp) | Coverage |
| --- | --- | --- | --- | --- |
| Female | *pair-end* | 150 | ~350 | 38.7X |
| Pool of females | *pair-end* | 150 | ~350 | 35.5X |

The final assemblies were around 524-538 Mb, with the pool being the less fragmented (**Table 2**).

**Table 2.**
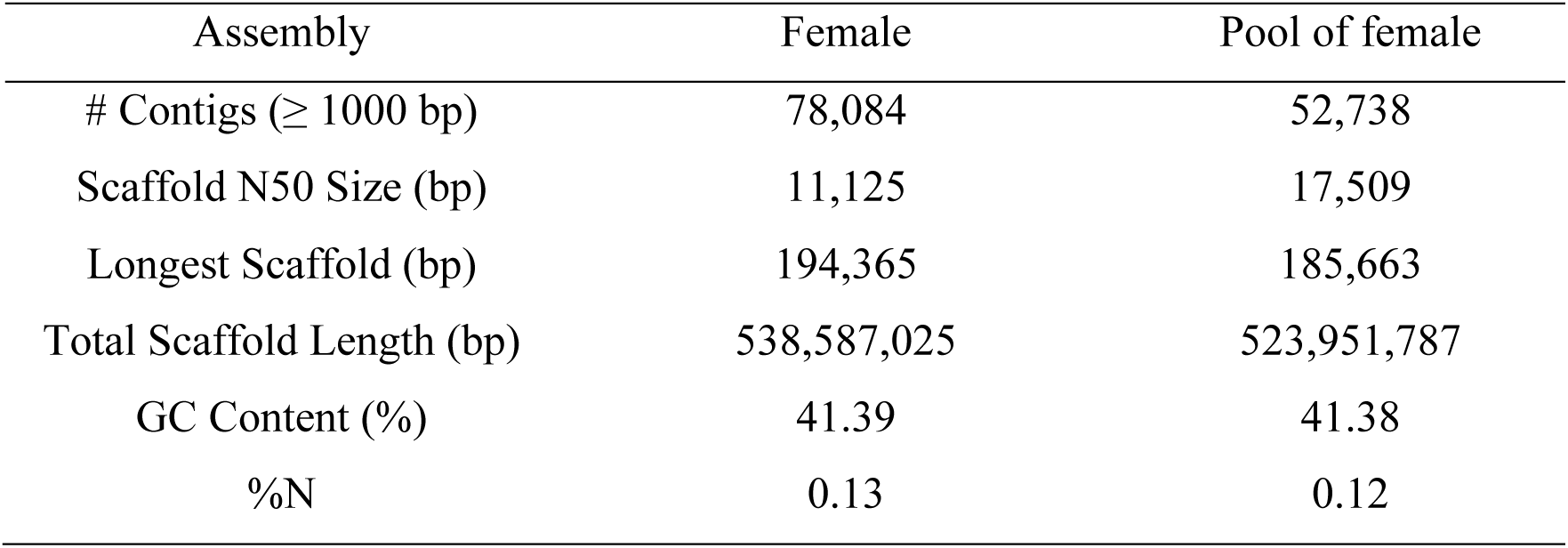
Black flounder assembly statistics

To verify annotation completeness, we checked for BUSCO (Waterhouse et al., 2018) orthologues in Actinopterygii DB, where 3459 (3252 complete and 198 fragmented, 94,7% of the selected database) were identified in the combined assemblies (for more information see **Table 3**).

**Table 3:** Gene product annotations results, GeneId, Exonerate, Infernal (with RFAM, DB, tRNAScan-SE and RNAmmer

|  | Female | Pool of female |
| --- | --- | --- |
| Proteins | 25,262 | 25,231 |
| KOG | 3541 | 3670 |
| Uncharacterized | ~4% | ~4% |
| tRNAs (all types) | 509 | 443 |
| rRNAs | 11 | 11 |
| snRNAs | 49 | 47 |
| Others ncRNAs | 971 | 889 |

### Transposable element identification and quantification

RepeatMasker showed large differences in the frequency of repetitive elements (REs) between fish species. **Fig. 1** compares TEs percentage and proportion among different fish genomes. In general, lineages with larger genome sizes had a higher frequency of REs (**Fig. 1**). In the same line, small genomes such as *T. nigroviridis* and Pleuronectiformes (especially *P. orbignyanus*) presented a very low frequency of REs. Among the Pleuronectiformes, the genome of *P. orbignyanus* has a lower proportion of DNA transposons and a higher proportion of long (LINEs) and short (SINEs) interspersed nuclear elements. We also studied the relationship between genome size with different genome elements, showing strong correlation with TEs percentage (r=0.847; p<0.001), total intron size (r=0.939; p<0.001) and total exon size (r=0.807; p<0.001) (**Fig. 2**).

**Figure 1.**
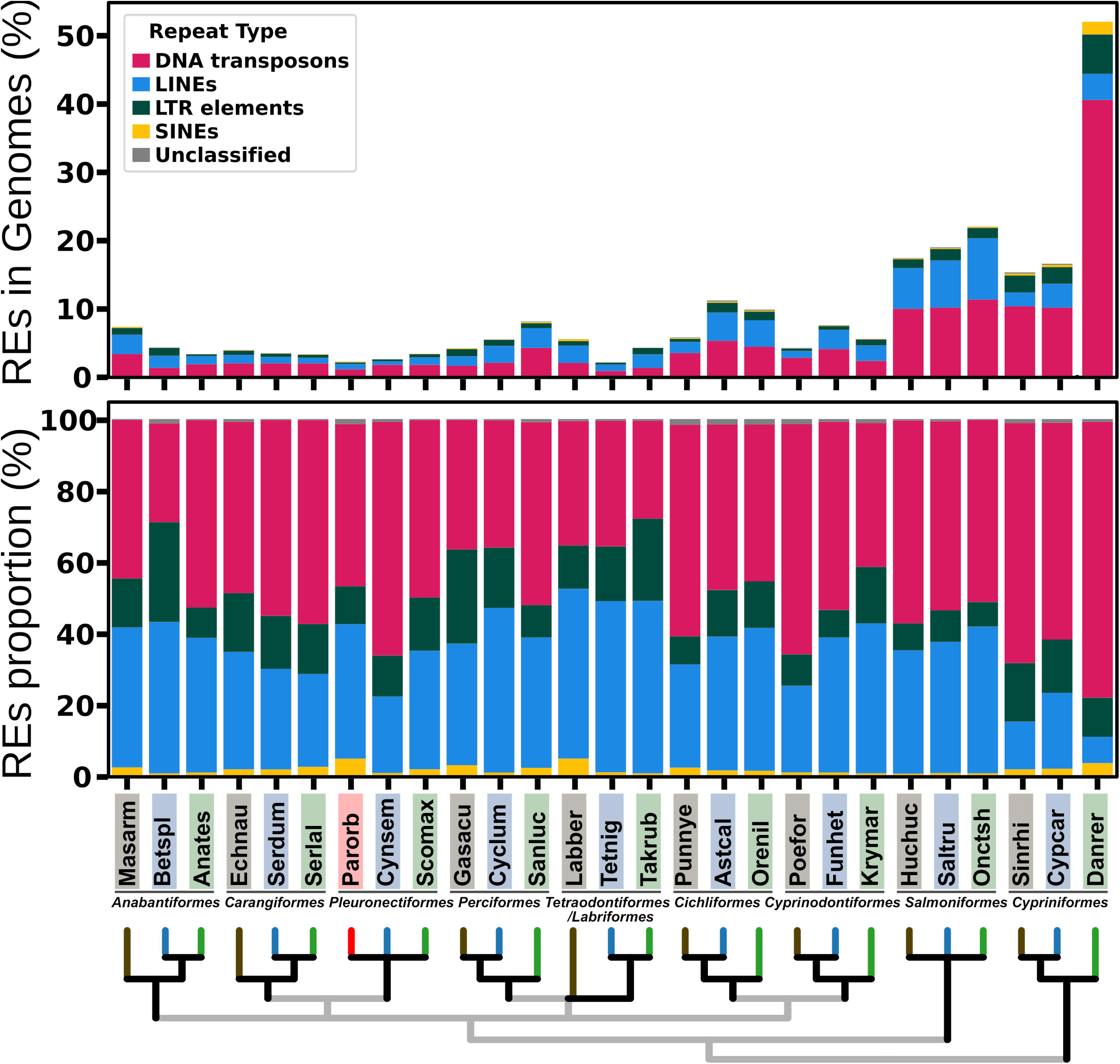
Contents of repetitive elements in fish genomes. Repetitive elements (REs) such as DNA transposons, long and short interspersed nuclear elements (LINEs and SINEs, respectively) and long terminal repeats (LTRs) are shown both as total percentage in genome (A) or percentage relative to repetitive elements content (B). A taxonomy tree for the species analyzed is shown below. Species: Anabantiformes (Anates: *Anabas testudineus*, Betspl: *Betta splendens*, Masarm: *Mastacembelus armatus*), Carangiformes (Echneu: *Echeneis naucrates*, Serdum: *Seriola dumerili*, Serlal: *Seriola lalandi dorsalis*), Cichliformes (Astcal: *Astatotilapia calliptera*, Orenil: *Oreochromis niloticus*, Punnye: *Pundamilia nyererei*), Cypriniformes (Cypcar: *Cyprinus carpio*, Danrer: *Danio rerio*, Sinrhi: *Sinocyclocheilus rhinocerous*), Cyprinodontiformes (Funhet: *Fundulus heteroclitus*, Krymar: *Kryptolebias marmoratus*, Poefor: *Poecilia formosa*), Perciformes (Cyclum: *Cyclopterus lumpus*, Gasacu: *Gasterosteus aculeatus*, Sanluc: *Sander lucioperca*), Pleuronectiformes (Cynsem: *Cynoglossus semilaevis*, Parorb: *Paralichthys orbygnanus*, Scomax: *Scophthalmus maximus*), Salmoniformes (Huchuc: *Hucho hucho*, Oncmyk: *Oncorhynchus mykiss*, Saltru: *Salmo trutta*), Labriformes (Labber: *Labrus bergylta*) and Tetraodontiformes (Takrub: *Takifugu rubripes*, Tetnig: *Tetraodon nigroviridis*).

**Figure 2.**
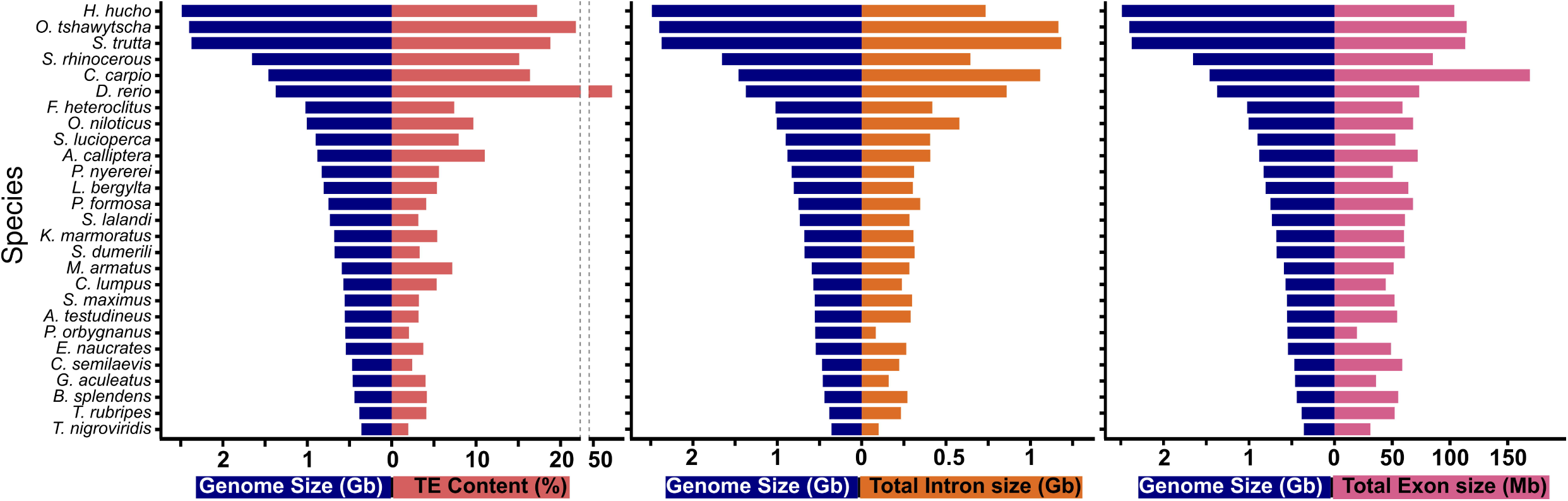
Correlation between genome size (Gb) and TE content (%) of fish.

### Gene family clustering

Gene families were clustered according to their similar structures into three different flatfish families; Scophthalmidae (*S. maximus*), Paralichthyidae (*P. orbignyanus and P. olivaceous*), Cynoglossidae (*C. semilaevis*), and *D. rerio* using OrthoMCL. A total of 11,870 putative specific genes among the five species and 1677 putative specific gene families were identified among the four flatfish species included in the analysis (**Fig. 3**).

**Figure 3.**
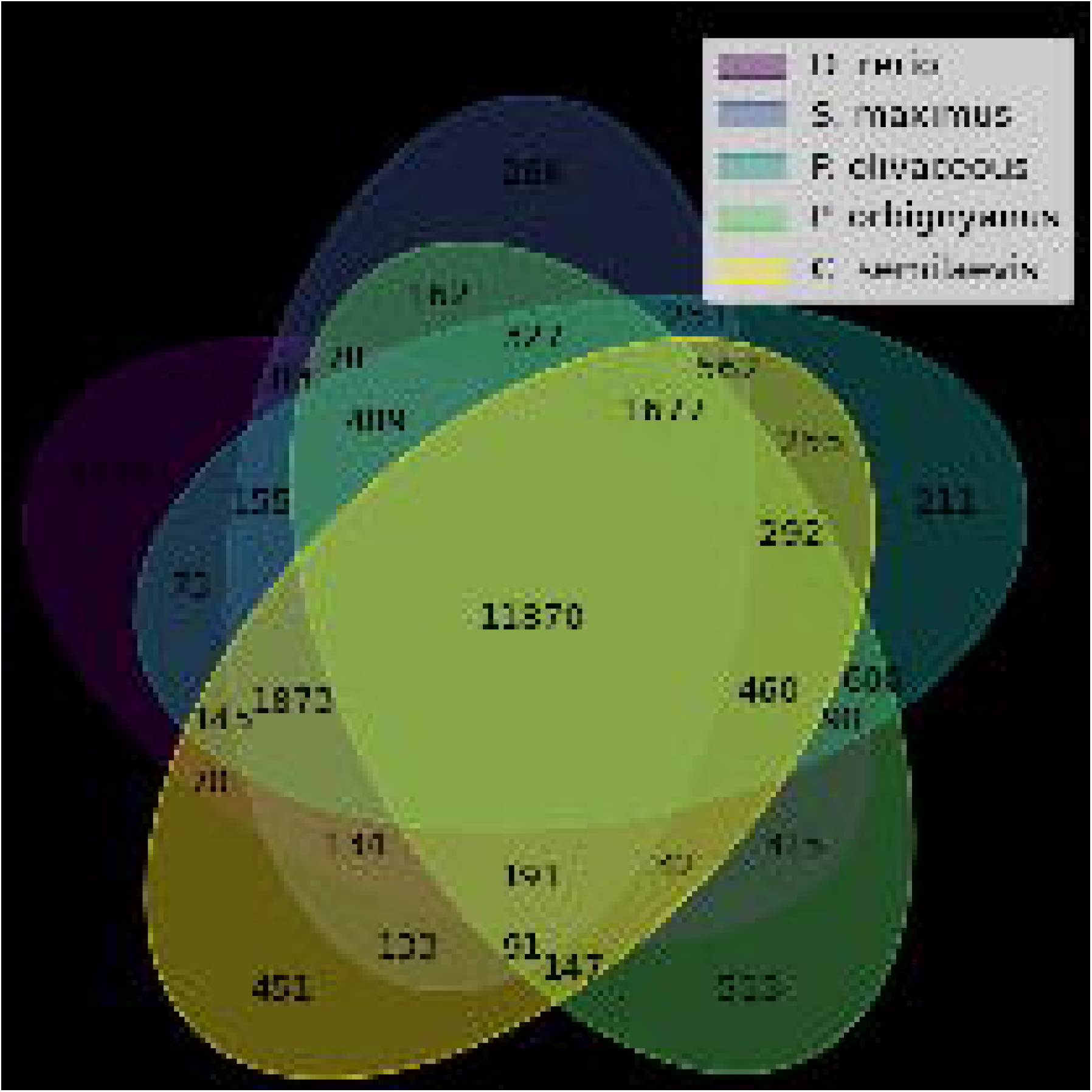
Venn diagram showing orthology in the three flatfish species available in Ensembl and zebrafish. Protein Orthologs were calculated using OrthoMCL (Li et al., 2003) with default parameters.

### C-value comparison between Pleuronectiformes and other fish orders

Using publicly available C-value data, we performed a comparative analysis of haploid genome sizes in fish (**Fig. 4, Supplementary Fig. S1**). We used data from 1504 species belonging to 73 orders (according to the NCBI taxonomy database). The estimated C-value of *P. orbignyanus* is 0.56 pg, which corresponds to a small fish genome size (5^th^ percentile = 0.61 pg). The average C-value for Pleuronectiformes is 0.734 pg, and the only order with a significantly lower average C-value is Tetraodontiformes (0.618 pg, *P* < 0.001). Gerreiformes, Ophidiiformes, Uranoscopiformes, Osmeriformes, and Chaetodontiformes also have lower average C-value than Pleuronectiformes, but statistical analysis did not reveal a significant difference. Eight of 49 species (16.3%) of Pleuronectiformes had C-values in the 5^th^ percentile range. In addition, 51.5% of Tetraodontiformes species (34/66) had C-values in the 5^th^ percentile, consistent with the fact that this order had the lowest mean C-value. Other orders represented in this 5^th^ percentile range are: Gerreiformes 2/4 (50%), Uranoscopiformes 2/4 (50%), Syngnathiformes 14/32 (43.75%), Ophidiiformes 2/5 (40%), Anabantiformes 5/28 (17.86%), Chaetodontiformes 2/15 (13.3%), Spariformes 5/39 (12.82%), Istiophoriformes 1/8 (12.5%), Carangiformes 3/26 (11.54%), Blenniiformes 5/44 (11.36%), Perciformes 8/95 (8.42%), Gobiiformes 3/40 (7.5%), Gadiformes 1/20 (5%), Acanthuriformes 1/23 (4.35%), Siluriformes 3/111 (2.70%), Lutjaniformes 1/37 (2.70%), Cypriniformes 8/394 (2.03%) and Centrarchiformes 1/51 (1.96%).

**Figure 4.**
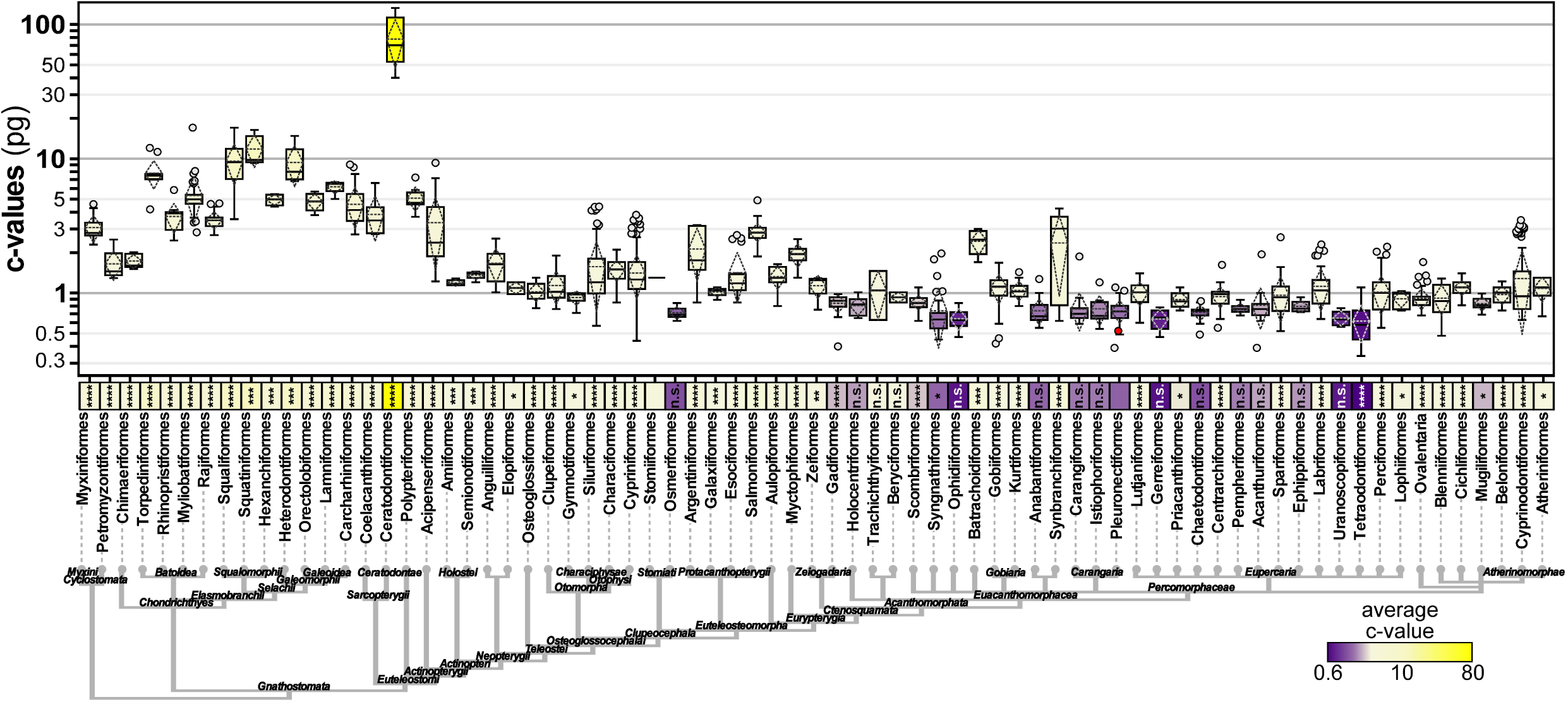
Haploid genome size across fish based on C-value. Boxplots representing C-values of 34 fish orders (top), organized by their taxonomic relationships (bottom). Y-axis represents C-values in log scale. For boxplots: horizontal bar, median; dashed lines, mean and standard deviation; filled circles, outliers; open circles, suspected outliers. C-value for *P. orbignyanus*, red colored circle. C-value means represented by boxes colored from yellow (highest) to purple (lowest). Analysis of unpaired t-test of *P. orbignyanus* versus each order are shown (****, P < 0.0001; ***, P < 0.001; **, P < 0.01; ns, not significant).

We additionally analyzed the C-values of all Pleuronectiformes species (**Fig. 5**). The C-value of *P. orbignyanus* is one of the smallest in the order (Pleuronectiformes 10^th^ percentile value = 0.55 pg), with three members of the family Paralichthyidae having lower C-values (*Pseudorhombus jenynsii* 0.54 pg, *Paralichthys dentatus* 0.53 pg, and *Pseudorhombus arsius* 0.49 pg). Finally, two other species from different families had smaller C-values: *Rhombosolea tapirina* 0.55 pg (Rhombosoleidae family), and *Pleuronectes platessa* 0.39 pg (Pleuronectidae family).

**Figure 5.**
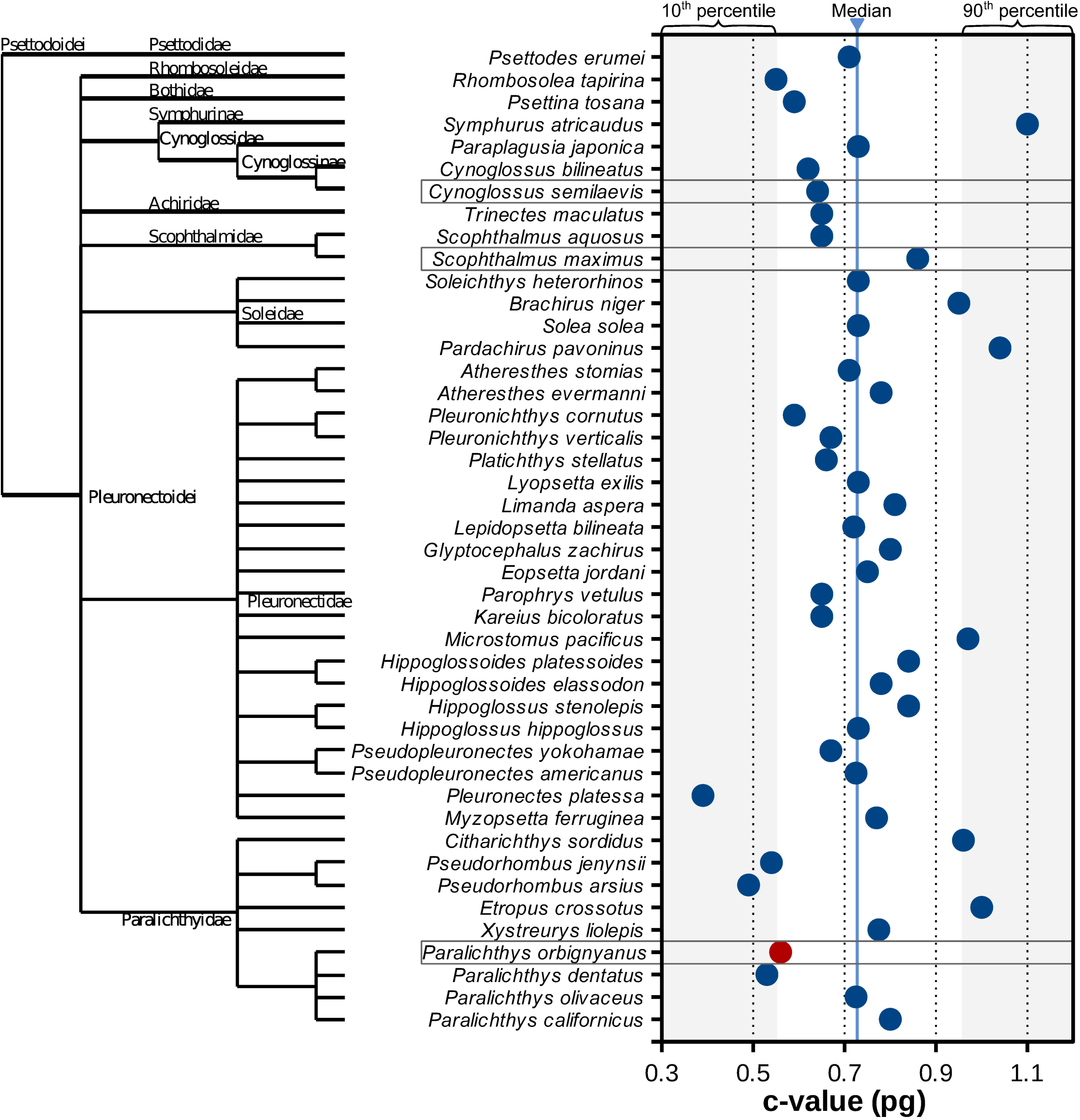
Pleuronectiformes species C-values. The median and the 10th and 90th percentile values are shown. C-value for *P. orbignyanus* shown in red. Other two species (*C. semilaevis* and *S. maximus*) with sequenced genomes are also highlighted. Taxonomy from NCBI’s Common Tree for Pleuronectiformes shown on the left.

Taken together, these results indicate that low genome size is a trait of Pleuronectiformes, and particularly in species from Paralichthyidae (including *P. orbignyanus*).

### Whole genome scale analysis of gene features size in teleost fish

To find a possible explanation for the relatively small size of the *P. orbignyanus* genome, we investigated whether this may be associated with major genetic feature sizes, e.g., smaller intron sizes. To this end, we determined the coordinates of all exons and introns for each gene model in the genome of *P. orbignyanus* and other twenty fish species. Using these data, we determined the sizes of all exons and introns, and inferred the size of the genes (exons sizes + introns sizes) and the size of the transcripts (exons sizes). Since it is possible to identify more than one gene model per locus, we also determined the number of exons (and introns) per gene model (**Fig. 6**). Remarkable, there is a reduction of the overall gene and transcript size in *P. orbignyanus*, compared to every other species analyzed here (**Fig. 6A** and **6B**), including species from Perciformes (*G. aculeatus*) and Tetraodontiformes (*T. nigroviridis*), which also show in general low C-values. Both gene and transcript size distributions are shifted to lower sizes in *P. orbignyanus*, observed in quantile analysis (**Supplementary Fig. S2** and **Supplementary Table Ss2-3**). Interestingly, the shift towards smaller sizes in *P. orbignyanus* compared to *T. nigroviridis* is more evident for transcript sizes. To discard an effect of differential untranslated regions sizes, we confirmed a shift towards smaller protein sizes in *P. orbignyanus* (**Supplementary Fig. S3**). In addition, we found that the number of exons and introns per gene model (**Fig. 6C**) in flounder is the lowest in the genomes analyzed (5.88 exons per model). Note that the total number of genes in *P. orbignyanus* (21531) is comparable to other species in Pleuronectiformes (21811 bp in *C. semilaevis* and 21447 in *S. maximus*). However, the transcript number is remarkably lower in *P. orbignyanus* compared to *C. semilaevis* and *S. maximus* (25262, 34435 and 35543, respectively) (**Fig. 6D** and **Supplementary Table Ss2-3**).

**Figure 6.**
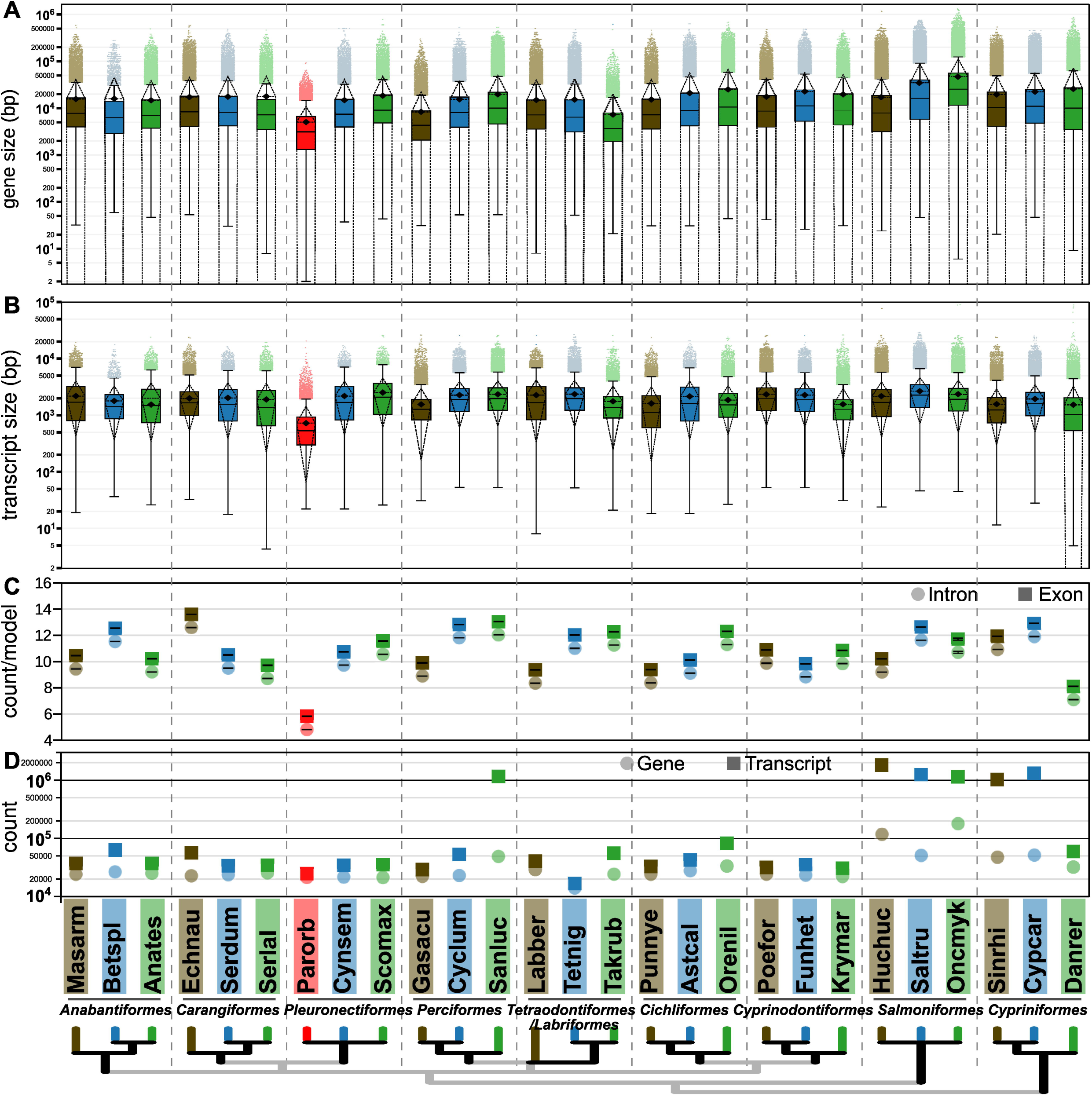
Boxplots representing whole genome distribution of genomic loci (A) and transcript models (B) sizes in fish. Average represented by diamonds, and standard deviation in dotted lines. (C) count of exons and introns in whole genomes. Average and SEM is shown. Six letter code for the species analyzed as described in Fig. 1.

Next, we studied the introns and exons size distribution at whole genome scale in fish genomes. Generally, we found that differences between average and median are smaller for exon sizes when compared to intron sizes, suggesting a more compact distribution of the former. Note also that tails towards higher feature size are larger for introns, suggesting that variability in intron size is more permissive towards larger introns rather than larger exons (**Fig. 7**).

**Figure 7.**
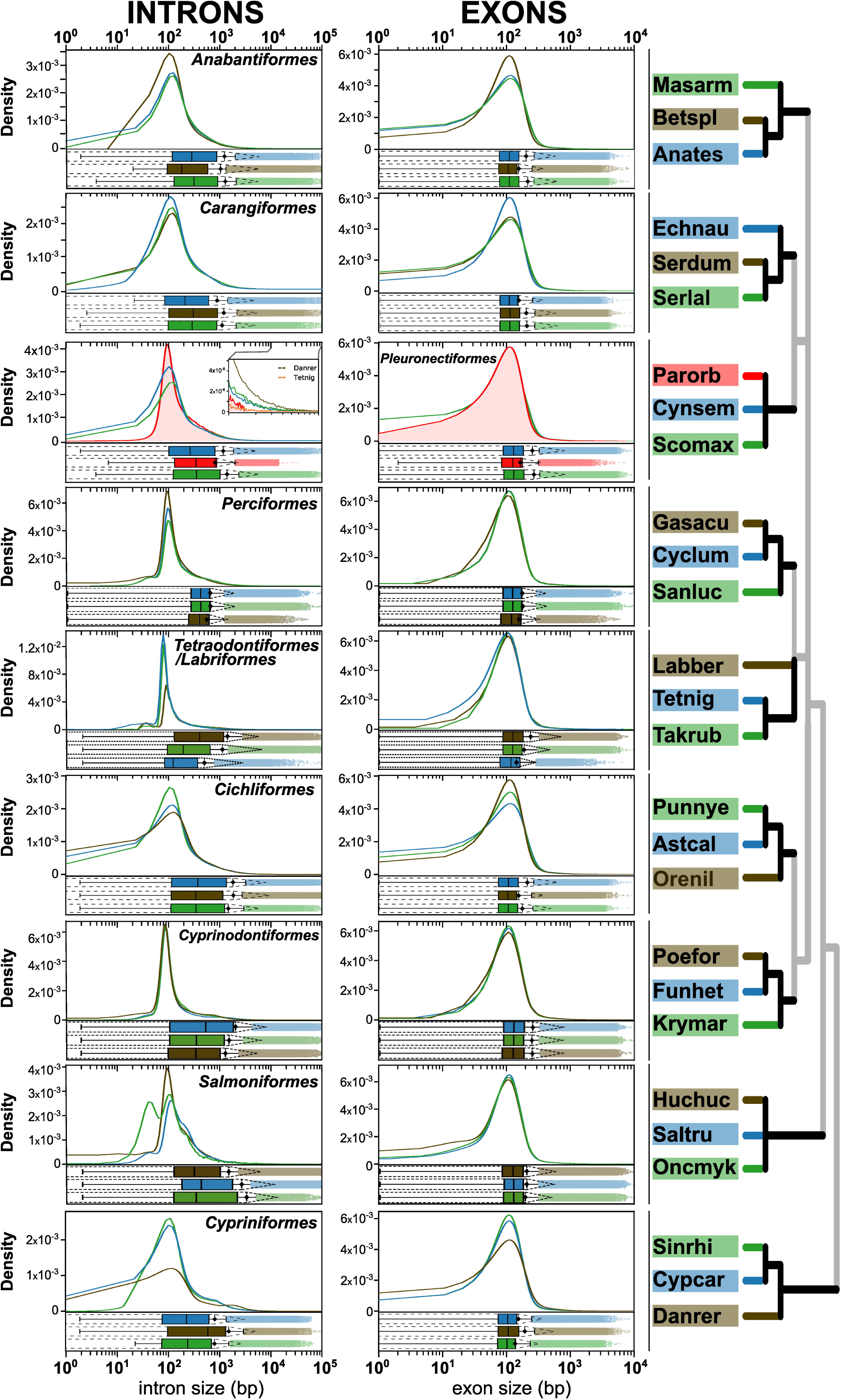
Size distribution of whole genome set of introns (left) and exons (right) for 27 species in 10 fish orders. In x-axis the size in bp is shown (log10 scale). KDE plots represent probability distribution of sizes (y-axis). Below each KDE plot, the boxplots represent the median (line) and quartiles 25 and 75%, whereas whiskers represent the upper and lower bounds. Mean (◆) ± s.d (dashed lines) are shown. Outliers set is shown with dots. Six letter code for the species analyzed as described in Fig. 1.

In general, we did not find major differences in the distribution of exon sizes between species. Median distribution values ranged from 119 bp (*T. nigroviridis*) to 132 bp (*F. heteroclitus*), while interquantile values 75^th^-25^th^ difference (IQR) span from 84 bp to 105 bp (*S. rhinocerous* and *P. formosa*, respectively). Among Pleuronectiformes, *P. orbignyanus* (median size value = 127 bp, IQR = 95 bp) has a smaller number of large exons, with the shift occurring more evidently at the 80^th^ quantile (**Fig. 7**, **Supplementary Fig. S4** and **Supplementary Table Ss4-5**). This shift in large exon size is also observed in species with small genome size, such as *T. nigroviridis* and *G. aculeatus*.

Greater variation among species is observed in intron sizes (**Fig. 7**, **Supplementary Fig. S5** and **Supplementary Table Ss4-5**). Median size values range from 121 bp (*T. nigroviridis*) to 1139 bp (*D. rerio*), and IQR values are highly variable, ranging from 286 in *T. nigroviridis* to 2602 in *D. rerio*. The largest difference in Pleuronectiformes is observed for large introns, with very large introns (>10000 bp) being extremely rare in *P. orbignyanus* (**Fig. 7**, see insert). The 99^th^ quantile for this species is 7311 bp, whereas in other Pleuronectiformes it is 16966 bp (*C. semilaevis*) and 19470 bp (*S. maximus*). The 99^th^ quantile in other orders is highly variable: in *T. nigroviridis*, the value of the 99^th^ quantile is 6002 bp (which is the lowest value) and increasing to 40237 bp in *D. rerio*.

## DISCUSSION

Fishes are the largest group of vertebrates, with more than 35,000 species (Fricke et al., 2020). About 20 years ago, the first complete genome of the Japanese pufferfish, *Takifugu rubripes*, was sequenced (Aparicio et al., 2002). Since then, the number of fish species sequences has greatly increased, largely due to advances in sequencing technologies and assembly algorithms (Ravi & Venkatesh, 2018). According to NCBI, nearly 3% (915) of all fish genomes have been sequenced as of August 2022. However, these numbers will soon be outdated due to large-scale sequencing of fish species genomes. For example, the 10,000 Fish Genomes Project (Fish10K) (Fan et al., 2020) aims to sequence and obtain reference genomes from representative fish species and has recently begun using long-read sequencing and Hi-C technology for at least one representative species from all families to obtain better quality reference genomes. All the information obtained from Fish10K, together with genome sequences from other laboratories (such as the one presented in this study), will help improve breeding programmes and promote sustainable aquaculture in the future, as well as perform genome editing and genomic selection (Lu & Luo, 2020).

Using next-generation sequencing (NGS) technologies, we were able to sequence, assemble, and annotate the black flounder genome. The black flounder genome was sequenced with high coverage and 25,231 protein-coding genes were identified. This is slightly more than the 21,787 protein-coding genes discovered in Japanese flounder (Shao et al., 2017). The estimated genome size of black flounder is ∼538 Mbp, similar to the genome size of other flatfish species: Japanese flounder (534 Mb) (Shao et al., 2017), Turbot (568 Mb) (Figueras et al., 2016), Spotted halibut (556 Mb) (Zhao et al., 2021), ∼15% bigger than the tongue sole genome (477 Mb) (Chen et al., 2014), and ∼11% smaller compared to the Senegalese sole genome (∼612 Mb) (Manchado et al., 2016). It is recommended to use multiple complementary methods to assess the genome quality (Gurevich et al., 2013). In this study, we assessed the N50 sizes of contigs and scaffolds and the completeness of the genome using KOG and BUSCO (Benchmarking Universal Single Copy Orthologs). The results of quantitative measurements to determine assembly completeness using BUSCO showed a high percentage of conserved orthologs in turbot (Xu et al., 2020). High BUSCO values (94.7 %) were observed in Actinopterygii using the combined assemblies, to compensate for the lack of long reads. Future transcriptome and long reads sequencing studies are therefore required to provide a comparative, more comprehensive and quantitative overview of the level of completeness achieved.

**Figure 3** shows an overlap of a total of 1677 putative orthologous groups exclusive for the three different flatfish families (Scophthalmidae, Cynoglossidae, and Paralichthyidae). It is important to note that a recent comparative genome analysis has examined the origins of flatfish body structure from an evolutionary perspective using the genomes of eight new species from fourteen families of Pleuronectiformes (Lü et al., 2021). This study will help to perform new comparative analyses in the future, including South American black flounder, once the annotated genomes become publicly available.

In this study, we performed a comparative analysis of 27 flatfish species in 10 orders to understand the small size of the black flounder genome. Genome size varies from very small genomes as in *Tetraodon nigroviridis* (∼350 Mb) (Neafsey and Palumbi, 2003) to large genomes as in *Salmo salar* (2967 Mb) (Yuan et al., 2018). As expected, the comparative analysis revealed that black flounder has one of the smallest genomes among the compared species, which is consistent with the results of previous studies in which flatfish genome sizes were among the smallest of all teleosts (Figueras et al., 2016; Xu et al., 2020). This observation is also consistent with the analysis of fish C-values (**Figures 4** and **5**). Indeed, the genomes of *B. splendens* (Anabantiformes), *G. aculeatus* (Perciformes, suborder Gasterosteiformes), *C. semilaevis* (Pleuronectiformes), *T. rubripes,* and *T. nigroviridis* (Tetraodontiformes) in the selected data set belong to groups with typically small genome size according to the average C-value (**Figure 4**). In addition, several C-values of Pleuronectioformes are within the 10th percentile of the distribution, especially in the families Paralichthyidae (including black flounder), Rhombosoleidae and Pleuronectidae. The large genome size in salmonids could be explained by a specific round (called 4R) whole-gene duplication event in this lineage (Lien et al., 2016). Although genome size diversity in teleosts could be the cause of the tremendous diversity of morphology, ecology, and behavior in this group (Volff, 2005), the origin of this dispersion is still unknown.

Genome size is not only affected by the teleost-specific rounds of whole-genome duplications (3R and 4R) (Meyer and Van de Peer, 2005). There are also other important events that seem to be involved, such as changes in the proportion of REs in genomes (Yun et al., 2018). Most of these REs come from specific sequences that replicate and move through the genome, called transposable elements (TEs) (Bourque, 2009). In animals, there is a correlation between genome size and the proportion of TEs (Canapa et al., 2015). Variation in genome size has been shown to be related to the amount of repetitive DNA in eukaryotic species (Kidwell, 2002), and this relationship has been clearly demonstrated in teleost fishes (Chalopin et al., 2015). The abundance of TEs varies with genome size and position in the fish tree of life (Shao et al., 2019). In this work, we compare the proportion of repetitive elements (REs): DNA transposons, long and short interspersed nuclear elements (LINEs and SINEs, respectively), and long terminal repeats (LTRs) in 27 fish species. In black flounder and the order of Pleuronectiformes in general, the proportion of all REs was low, which is probably one of the reasons for the small size of the black flounder genome. The low proportion of all REs contrasts with the high levels observed in zebrafish, as also noted in other studies (Lü et al., 2021). Thus, black flounder is consistent with the compact genomes observed in other flatfish genomes, with ∼5% TEs, slightly more than the 3% in pufferfish (Aparicio et al., 2002).

The number of exons and introns could also explain the genome size in teleost fish. To this end, the coordinates of all exons and introns in the genomes of twenty-one fish species, including *P. orbignyanus,* were determined to determine whether the small size of the *P. orbignyanus* genome was due to this. Results showed that *P. orbignyanus* has smaller gene and transcript sizes compared to all fish species studied, including groups with small genomes such as the Tetraodontiformes (Binerd et al., 2001) and some Perciformes (*e.g*., *G. aculeatus*) (Reid et al., 2021). Flow cytometric analyses revealed that in four pufferfish species of the family Tetraodontidae, genome size varies between 0.38 and 0.82 pg, whereas the sister family Diodontidae is larger (0.8-1 pg), likely due to DNA loss during divergence of the two families during evolution (Noleto et al., 2009). This is consistent with our study, which showed that all pleuronectiformes generally have a small genome size (0.3-1.1 pg).

In eukaryotes, intronic DNA is the major component of genes and genomes and plays a key role in gene regulation, and intron size is important from an evolutionary perspective (Zhang and Edwards, 2012). In teleosts, genome size and intron size are closely related, with intron size reflecting genome size (Jakt et al., 2022). To further investigate the origin of the small genome size of black flounder, we examined the distribution of total genome size of introns and exons for 27 species in 10 fish orders. Our analyses show that the mean and median exon sizes are smaller compared to intron sizes, suggesting a more compact distribution, as observed in other Pleuronectiformes species (Robledo et al., 2017). As observed in previous studies, no differences in exon sizes were detected between species (Li et al., 2017). Knowledge of exon *loci* is important for phylogenomic analyses (Hughes et al., 2021).

Based on the analyzed features, we concluded that the most important components that could be responsible for the reduced flounder genome are (i) the low frequency of repetitive elements, (ii) the reduced transcript size, which is most likely due to a lower number of exons in the whole genome and probably has a lower impact, (iii) the lower number of very large introns. The last two components (ii and iii) have a lower value than in other species with similar C-values, suggesting that this may be a novel genome reduction strategy.

In summary, in this study we generated a genome assembly of black flounder (*Paralichthys orbignyanus*). The results show a reduced genome size of flounder, and we have demonstrated that this is related to reduced transcript size due to a reduction in exon and intron sizes. Finally, our study also serves to investigate a strategy of genome reduction in teleost fishes.

## ACKNOWLEDGEMENTS

We thank the Laboratorio de Maricultura del Instituto Nacional de Investigación y Desarrollo Pesquero (INIDEP), Argentina, and E. Aristizabal for kindly providing tissue samples. The authors thank J.J. Boccanfuso and E. Ricci for their help in collecting fish samples or transporting them to Europe; M.B. Gomes Pardo for DNA extraction; and Prof. P. Martinez Portela for helpful comments on this manuscript. We would like to acknowledge the support of the Centro de Supercomputación de Galicia (CESGA) in the completion of this work. Finally, this article is dedicated to our colleague and friend Nicolás Stocchi, who recently passed away.

## AVAILABILITY OF SUPPORTING DATA

This sequencing data has been deposited at the National Center for Biotechnology Information (NCBI) where the Bioproject Accession ID is PRJEB36690.

## Supplementary Fig S1

Correlation plots for C-values (as per Animal Genome size database or estimated from sequenced genome, when available, in x-axis) vs feature count (top, log scale), feature median size (center, bp) and feature 99^th^ percentile size (bottom, bp). Features analyzed are Genes, Transcripts, Intron and Exons (in columns from left to right). Data corresponding to *P. orbignyanus* highlighted in red.

## Supplementary Fig S2 Quantile_gene-transcript

For each species, gene (left) and transcript (right) size distribution *n*^th^ percentiles (*n* 1, 5, 10, 20, 25, 30, 40, 50, 60, 75, 80, 90, 95 and 99) were computed (shown in y-axis size, bp in log scale). *P. orbignyanus* data (red) is shown in all plots for comparison. Six letter code for the species analyzed as described in Fig. 1.

## Supplementary Fig S3 boxplot_protein

A. Complete fish proteome size distribution. Horizontal bar shows median, whereas dotted lines represent standard deviation, and diamonds represent mean value. Outliers are shown as filled circles. Six letter code for the species analyzed as described in Fig. 1. B. Kernel density estimation plot of protein size distribution in Pleuronectiformes (Cynsem, Scomax and Parorb), plus representatives from Tetraodontiformes (Tetnig) and Cypriniformes (Danrer).

## Supplementary Fig S4 Quantile_exon_intron

For each species, exon (left) and intron (right) size distribution *n*^th^ percentiles (*n* 1, 5, 10, 20, 25, 30, 40, 50, 60, 75, 80, 90, 95 and 99) were computed (shown in y-axis size, bp in log scale). *P. orbignyanus* data (red) is shown in all plots for comparison. Six letter code for the species analyzed as described in Fig. 1.

## REFERENCES

Aparicio, S., Chapman, J., Stupka, E., Putnam, N., Chia, J.M., Dehal, P., Christoffels, A., Rash, S., Hoon, S., Smit, A., Gelpke, M.D.S., Roach, J., Oh, T., Ho, I.Y., Wong, M., Detter, C., Verhoef, F., Predki, P., Tay, A., Lucas, S., Richardson, P., Smith, S.F., Clark, M.S., Edwards, Y.J.K., Doggett, N., Zharkikh, A., Tavtigian, S.V., Pruss, D., Barnstead, M., Evans, C., Baden, H., Powell, J., Glusman, G., Rowen, L., Hood, L., Tan, Y.H., Elgar, G., Hawkins, T., Venkatesh, B., Rokhsar, D., Brenner, S., 2002. Whole-genome shotgun assembly and analysis of the genome of Fugu rubripes. Science, 297, 1301–1310. https://doi.org/10.1126/science.1072104

Bambill, G.A., Masakazu, O., Radonic, M., López, A.V., Müller, M.I., Boccanfuso, J.J., Bianca, F.A., 2006. Broodstock management and induced spawning of flounder (Paralichthys orbignyanus) (Valenciennes, 1893) under a closed recirculated system. Revista de Biología Marina y Oceanográfica, 41, 45–55. http://dx.doi.org/10.4067/S0718-19572006000100007

Basu, S., Hadzhiev, Y., Petrosino, G. Nepal, C., Gehrig, J., Armant, O., Ferg, M., Strahle, U., Sanges, R., Müller, F., 2016. The *Tetraodon nigroviridis* reference transcriptome: developmental transition, length retention and microsynteny of long non-coding RNAs in a compact vertebrate genome. Sci Rep, 6, 33210. https://doi.org/10.1038/srep33210

Bao, W, Jurka, M.G., Kapitonov, V.V., Jurka, J., 2009. New superfamilies of eukaryotic DNA transposons and their internal divisions. Molecular Biology and Evolution 26(5), 983–293. https://doi.org/10.1093/molbev/msp013

Blanco, E.; Abril, J.F., 2009. Computational gene annotation in new genome assemblies using GeneID. Methods Mol Biol, 537, 243–261. https://doi.org/10.1007/978-1-59745-251-9_12

Blanquer, A. (1990). Phylogéographie intraspécifique d’un poisson marin, le flet Platichthys flesus L. (Heterosomata): Polymorphisme des marqueurs nucléaires et mitochondriaux [Thesis, Montpellier 2]. In Http://www.theses.fr. http://www.theses.fr/1990MON20015

Brainers, E.L., Slutz, S.S., Hall, E.K., Phillips, W.R., 2001. Patterns of genome size evolution in tetraodontiform fishes. Evolution 55(11), 2363-2368. https://doi.org/10.1111/j.0014-3820.2001.tb00750

Brenner, S., Elgar, G., Sandford, R., Macrae, A., Venkatesh, B., Aparicio, S., 1993. Characterization of the pufferfish (Fugu) genome as a compact model vertebrate genome. Nature, 366(6452), 265–8. https://doi.org/10.1038/366265a0

Boccanfuso, J.J., Aristizabal, A.E.O., Berrueta, M., 2019. Improvement of natural spawning of black flounder, Paralichthys orbignyanus (Valenciennes, 1839) by photothermal and salinity conditioning in recirculating aquaculture system. Aquaculture, 502, 134–141. https://doi.org/10.1016/j.aquaculture.2018.12.034

Boetzer, M., Henkel, C.V., Jansen, H.J., Butler, D., Pirovano, W., 2011. Scaffolding pre-assembled contigs using SSPACE. Bioinformatics, 27, 578–579. https://doi.org/10.1093/bioinformatics/btq683

Boetzer, M., Pirovano, W., 2012. Toward almost closed genomes with GapFiller. Genome Biol, 13: R56. https://doi.org/10.1186/gb-2012-13-6-r56

Bolger, A.M., Lohse, M., Usadel, B., 2014. Trimmomatic: A flexible trimmer for Illumina sequence data. Bioinformatics, 30, 2114–2120. https://doi.org/10.1093/bi-oinformatics/btu170

Bourque, G., 2009. Transposable elements in gene regulation and in the evolution of vertebrate genomes. Current Opinion in Genetics & Development, 19(6), 607–612. https://doi.org/10.1016/j.gde.2009.10.013

Brainerd, E.L., Slutz, S.S., Hall, E.K., Phillis, R.W. 2001. Patterns of genome size evolution in Tetraodontiform fishes. Evolution, 55, 2363–2368. https://doi.org/10.1111/j.0014-3820.2001.tb00750.x

Canapa, A., Barucca, M., Biscotti, M.A., Forconi, M., Olmo, E., 2015. Transposons, Genome Size, and Evolutionary Insights in Animals. Cytogenetic and genome research, 147(4), 217–239. https://doi.org/10.1159/000444429

Chalopin, D., Naville, M., Plard, F., Galiana, D., Volff, J.N., 2015. Comparative analysis of transposable elements highlights mobilome diversity and evolution in vertebrates. Genome Biology and Evolution, 7(2), 567–580. https://doi.org/10.1093/gbe/evv005

Cerdà, J., Douglas, S., Reith, M., 2010. Genomic Resources for Flatfish Research and Their Applications. Journal of Fish Biology 77(5), 1045–1070. https://doi.org/10.1111/j.1095-8649.2010.02695.x

Chen, S., Zhang, G., Shao, C., Huang, Q., Liu, G., Zhang, P., Song, W., An, N., Chalopin, D., Volff, J.N., Hong, Y., Li, Q., Sha, Z., Zhou, H., Xie, M., Yu, Q., Liu, Y., Xiang, H., Wang, N., Wu, K., Yang, C., Zhou, Q., Liao, X., Yang, L., Hu, Q., Zhang, J., Meng, L., Jin, L., Tian, Y., Lian, J., Yang, J., Miao, G., Liu, S., Liang, Z., Yan, F., Li, Y., Sun, B., Zhang, H., Zhang, J., Zhu, Y., Du, M., Zhao, Y., Schartl, M., Tang, Q., Wang, J., 2014. Whole-genome sequence of a flatfish provides insights into ZW sex chromosome evolution and adaptation to a benthic lifestyle. Nature Genetics 46, 253–260. https://doi.org/10.1038/ng.2890

Díaz De Astarloa, J.M., Munroe, T.A., 1998. Systematics, distribution and ecology of commercially important paralichthyd flounders occuring in Argentina-Uruguayan waters (Paralichthys, Paralichthydae). An overview. J. Sea Res. 39, 1**–**9. https://doi.org/10.1016/S1385-1101(97)00010-5

Díaz de Astarloa, J.M., 2002. A review of the flatfish fisheries of the south Atlantic Ocean. Revista de Biología Marina y Oceanografía, 37(2), 113-125. http://dx.doi.org/10.4067/S0718-19572002000200001

Doležel, J., Bartoš, J., Voglmayr, H., Greilhuber, J., 2003. Nuclear DNA content and genome size of trout and human. Cytometry A 51, 127–128. https://doi.org/10.1002/cyto.a.10013

Einfeldt, A.L., Kess, T., Messmer, A., Duffy, S., Wringe, B.F., Fisher, J., den Heyer, C., Bradbury, I.R., Ruzzante, D.E., Bentzen, P., (2021), Chromosome level reference of Atlantic halibut *Hippoglossus hippoglossus* provides insight into the evolution of sexual determination systems. Mol Ecol Resour, 21, 1686–1696. https://doi.org/10.1111/1755-0998.13369

Fan, G., Song, Y., Yang, L., Huang, X., Zhang, S., Zhang, M., Yang, X., Chang, Y., Zhang, H., Li, Y., Liu, S., Yu, L., Chu, J., Seim, I., Feng, C., Near, T.J., Wing, R.A., Wang, W., Wang, K., Wang, J., Xu, X., Yang, H., Liu, X., Chen, N., He, S., 2020. Initial data release and announcement of the 10,000 Fish Genomes Project (Fish10K), GigaScience 9, 8. https://doi.org/10.1093/gigasci-ence/giaa080

Ferreira, D.C., Porto-Foresti, F., Oliveira, C., Foresti, F., 2011. Transposable elements as a potential source for understanding the fish genome. Mobile genetic elements, 1(2), 112–117. https://doi.org/10.4161/mge.1.2.16731

Figueras, A., Robledo, D., Corvelo, A., Hermida, M., Pereiro, P., Rubiolo, J.A., Gómez-Garrido, J., Carreté, L., Bello, X., Gut, M., Gut, I.G., Marcet-Houben, M., Forn-Cuní, G., Galán, B., García, J.L., Abal-Fabeiro, J.L., Pardo, B.G., Taboada, X., Fernández, C., Vlasova, A., Hermoso-Pulido, A., Guigó, R., Álvarez-Dios, J.A., Gómez-Tato, A., Viñas, A., Maside, X., Gabaldón, T., Novoa, B., Bouza, C., Alioto, T., Martínez, P., 2016. Whole genome sequencing of turbot (*Scophthalmus maximus*; Pleuronectiformes): a fish adapted to demersal life. DNA Research 23, 181–192. https://doi.org/10.1093/dnares/dsw007

Fricke, R., Eschmeyer, W.N., van der Laan, R. (Eds.), 2020. Eschmeyeŕs Catalog of Fishes: Genera. Species, References http://researcharchive.calacademy.org/re-search/ichthyology/catalog/fishcatmain.asp

Gao, B., Shen, D., Xue, S., Chen, C., Cui, H., Song, C., 2016. The contribution of transposable elements to size variations between four teleost genomes. Mob DNA, 7:4. https://doi.org/10.1186/s13100-016-0059-7

Gregory, T.R., 2020. Animal Genome Size Database. http://www.genomesize.com

Griffiths-Jones, S., Bateman, A., Marshall, M., Khanna, A., Eddy, S.R., 2003. Rfam: an RNA family database. Nucleic Acids Res, 31, 439–441. https://doi.org/10.1093/nar/gkg006

Guerrero-Cózar, I., Gomez-Garrido, J., Berbel, C., Martinez-Blanch, J.F., Alioto, T., Claros, M.G., Gagnaire, P.A., Manchado, M. 2021. Chromosome anchoring in Senegalese sole (*Solea senegalensis*) reveals sex-associated markers and genome rearrangements in flatfish. Sci Rep 11, 13460. https://doi.org/10.1038/s41598-021-92601-5

Gurevich, A., Saveliev, V., Vyahhi, N., Tesler, G., 2013. QUAST: Quality assessment tool for genome assemblies. Bioinformatics 29, 1072–1075. https://doi.org/10.1093/bioinformatics/btt086

Hancock, J.M., 2004. Repeatmasker. *In*: Dictionary of Bioinformatics and Computational Biology. John Wiley & Sons, Ltd, Chichester, p. dob0206. https://doi.org/10.1002/9780471650126.dob0065.pub2

Hancock, J.M., Bishop, M.J., 2004. InterProScan. *In*: Dictionary of Bioinformatics and Computational Biology. John Wiley & Sons, Ltd, Chichester, p.206.

Hinegardner, R., 1968. Evolution if cellular DNA content in teleost fishes. American Naturalist 102, 517-523. https://doi.org/10.1086/282564

Hinegardner, R., Rosen, D.E., 1972. Cellular DNA content and the evolution of teleostean fishes. American Naturalist 106, 621-644. https://doi.org/10.1086/282801

Hubley, R., Finn, R.D., Clements, J., Eddy, S.R., Jones, T.A., Bao, W., Smit, A.F.A., Wheeler, T.J., 2016. The Dfam database of repetitive DNA families. Nucleic Acids Res, 44, 81–89. https://doi.org/10.1093/nar/gkv1272

Hughes, L.C., Ortí, G., Saad, H., Li, C., White, W.T., Baldwin, C.C., Crandall, K.A., Arcila, D., Betancur-R, R., 2021. Exon probe sets and bioinformatics pipelines for all levels of fish phylogenomics. Mol Ecol Resour. 21(3), 816–833. https://doi.org/10.1111/1755-0998.13287

Jakt, L.M., Dubin, A., Johansen, S.D., 2022. Intron size minimisation in teleosts. BMC Genomics 23, 628 (2022). https://doi.org/10.1186/s12864-022-08760-w

Kelley, D.R., Schatz, M.C., Salzberg, L.L., 2010. Quake: Quality-aware detection and correction of sequencing errors. Genome Biol, 11, R116. https://doi.org/10.1186/gb-2010-11-11-r116

Kidwell, M.G. 2002. Transposable elements and the evolution of genome size in eukaryotes. Genetica 115, 49–63. https://doi.org/10.1023/A:1016072014259

Lagesen, K., Hallin, P., Rødland, E.A., Staerfeldt, H.H., Rognes, T., Ussery, D.W., 2007. RNAmmer: consistent and rapid annotation of ribosomal RNA genes. Nucleic Acids Res. 35, 3100–3108. https://doi.org/10.1093/nar/gkm160

Li, L., Stoeckert, C.J.Jr., Roos, D.S., 2003. OrthoMCL: identification of ortholog groups for eukaryotic genomes. Genome research, 13(9), 2178–2189. https://doi.org/10.1101/gr.1224503

Li, Y., Xu, Y., Ma, Z., 2017. Comparative Analysis of the Exon-Intron Structure in Eukaryotic Genomes. Yangtze Medicine, 1, 50–64. https://doi.org/10.4236/ym.2017.11006

Lien, S., Koop, B.F., Sandve, S.R., Miller, J.R., Kent, M.P., Nome, T., Hvidsten, T.R., Leong, J.S., Minkley, D.R., Zimin, A., Grammes, F., Grove, H., Gjuvsland, A., Walenz, B., Hermansen, R.A., von Schalburg, K., Rondeau, E.B., Di Genova, A., Samy, J.K.A., Olav Vik, J., Vigeland, M.D., Caler, L., Grimholt, U., Jentoft, S., Inge Våge, D., de Jong, P., Moen, T., Baranski, M., Palti, Y., Smith, D.R., Yorke, J.A., Nederbragt, A.J., Tooming-Klunderud, A., Jakobsen, K.S., Jiang, X., Fan, D., Hu, Y., Liberles, D.A., Vidal, R., Iturra, P., Jones, S.J.M., Jonassen, I., Maass, A., Omholt, S.W., Davidson, W.S. 2016. The Atlantic salmon genome provides insights into rediploidization. Nature 533, 200–205. https://doi.org/10.1038/nature17164

López, A.V., Müller, M.I., Radonic, M., Bambill, G.A., Boccanfuso, J.J., Bianca, F.A., 2009. Larval culture technique and quality control in juveniles of flounder Paralichthys orbignyanus (Valenciennes, 1839) in Argentina. Spanish Journal of Agricultural Research, 75-82. http://dx.doi.org/10.5424/sjar/2009071-400

Lu, G., Luo, M., 2020. Genomes of major fishes in world fisheries and aquaculture: Status, application and perspective. Aquaculture and Fisheries, 5, 163–173. https://doi.org/10.1016/j.aaf.2020.05.004

Lu, J.G., Peatman, E., Wang, W.Q., Yang, Q., Abernathy, J., Wang, S.L., Kucuktas, H., Liu, Z.J., 2010. Alternative splicing in teleost fish genomes: same-species and cross-species analysis and comparisons. Mol. Genet. Genom. 283, 531–539. https://doi.org/10.1007/s00438-010-0538-3

Luckenbach, J.A., Borski, R.J., Daniels, H.V., Godwin, J., 2009. Sex determination in flatfishes: Mechanisms and environmental influences. Seminars in Cell & Developmental Biology, 20, 256–263, https://doi.org/10.1016/j.semcdb.2008.12.002

Luo, R., Liu, B., Xie, Y., Li, Z., Huang, W., Yuan, J., He, G., Chen, Y., Pan, Q., Liu, Y., Tang, J., Wu, G., Zhang, H., Shi, Y., Liu, Y., Yu, C., Wang, B., Lu, Y., Han, C., Cheung, D.W., Yiu, S., Peng, S., Zhu, X., Liu, G., Liao, X., Li, Y., Yang, H., Wang, J., Lam, T., Wang, J., 2012. SOAPdenovo2: an empirically improved memory-efficient short-read de novo assembler. Gigascience 4, 18. http://dx.doi.org/10.1186/2047-217X-1-18

Lü, Z., Gong, L., Ren, Y., Chen, Y., Wang, Z., Liu, L., Li, H., Chen, X., Li, Z., Luo, H., Jiang, H., Zeng, Y., Wang, Y., Wang, K., Zhang, C., Jiang, H., Wan, W., Qin, Y., Zhang, J., Zhu, L., Shi, W., He, S., Mao, B., Wang, W., Kong, X., Li, Y., 2021. Large-scale sequencing of flatfish genomes provides insights into the polyphyletic origin of their specialized body plan. Nat Genet. https://doi.org/10.1038/s41588-021-00836-9

Magnone, L., Bessonart, M., Rocamora, M., Gadea, J., Salhi, M., 2015. Diet estimation of *Paralichthys orbignyanus* in a coastal lagoon via quantitative fatty acid signature analysis. J Exp Mar Biol Ecol 462, 36–49. https://doi.org/10.1016/j.jembe.2014.10.008

Martínez, P., Robledo, D., Taboada, X., Blanco, A., Moser, M., Maroso, F., Hermida, M., Gómez-Tato, A., Álvarez-Blázquez, B., Cabaleiro, S., Piferrer, F., Bouza, C., Lien, S., Viñas, A., 2021. A genome-wide association study, supported by a new chromosome-level genome assembly, suggests sox2 as a main driver of the undifferentiatiated ZZ/ZW sex determination of turbot (*Scophthalmus maximus*). Genomics, 113, 1705–1718, https://doi.org/10.1016/j.ygeno.2021.04.007

Maroso, F., Hermida, M., Millán, A., Blanco, A., Saura, M., Fernández, A., Dalla Rovere, G., Bargelloni, L., Cabaleiro, S., Villanueva, B., Bouza, C., Martínez, P. 2018. Highly dense linkage maps from 31 full-sibling families of turbot (*Scophthalmus maximus*) provide insights into recombination patterns and chromosome rearrangements throughout a newly refined genome assembly. DNA Res. 25, 439–450. https://doi.org/10.1093/dnares/dsy015

Manchado, M., Planas, J.V., Cousin, X., Rebordinos, L., Claros, M.G., 2016. Current status in other finfish species: Description of current genomic resources for the gilthead seabream (*Sparus aurata*) and soles (*Solea senegalensis* and *Solea solea*), in *Genomics in Aquaculture*, eds S. Mackenzie and S. Jentoft (Amsterdam: Elsevier), 195–221. https://doi.org/10.1016/B978-0-12-801418-9.00008-1

Mank, J.E., Avise, J.C., 2006. Phylogenetic conservation of chromosome numbers in Actinopterygiian fishes. Genetica 127, 321–7. https://doi.org/10.1007/s10709-005-5248-0

Meyer, A., Van de Peer, Y., 2005. From 2R to 3R: evidence for a fish-specific genome duplication (FSGD). BioEssays, 27, 937–45. https://doi.org/10.1002/bies.20293

Nawrocki, E.P., Eddy, S.R., 2013. Infernal 1.1: 100-fold faster RNA homology searches. Bioinformatics. 29, 2933-2935 https://doi.org/10.1093/bioinformat-ics/btt509

Neafsey, D.E., Palumbi, S.R., 2003. Genome size evolution in pufferfish: a comparative analysis of diodontid and tetraodontid pufferfish genomes. Genome Res, 13(5), 821–30. http://www.genome.org/cgi/doi/10.1101/gr.841703.

Nelson, J.S., Grande, T.C., Wilson, M.V.H., 2016. Fishes of the World. 5th Edition, John Wiley and Sons, Hoboken. https://doi.org/10.1002/9781119174844

Noleto, R.B., de Souza Fonseca Guimarães, F., Paludo, K.S., Vicari, M.R., Artoni, R.F., Cestari, M.M., 2009. Genome size evaluation in Tetraodontiform fishes from the Neotropical region. Mar Biotechnol 11(6), 680–5. https://doi.org/10.1007/s10126-009-9215-0

Pundir, S., Martin, M.J., O’donovan, C., 2017. UniProt protein knowledgebase. Methods Mol Biol, 1558, 41–55. https://doi.org/10.1007/978-1-4939-6783-4_2

Radonic, M., Müller, M.I., López, A.V., Bambill, G.A., Spinedi, M.; Boccanfuso, J.J., 2007. Improvement on flounder Paralichthys orbignyanus (Valenciennes, 1839) controlled spawning in Argentina. Cien. Mar. 33 (2), 187–196. https://doi.org/10.7773/cm.v33i2.1021

Radonic, M., Macchi, G.J. 2009. Gonadal sex differentiation in cultured juvenile flounder, Paralichthys orbignyanus (Valenciennes, 1839). Journal of the World Aquaculture Society, 40, 129-133. https://doi.org/10.1111/j.1749-7345.2008.00229.x

Ravi, V., Venkatesh, B., 2018. The Divergent Genomes of Teleosts. Annu. Rev. Anim. Biosci, 6, 47–68. https://doi.org/10.1146/annurev-animal-030117-014821

Read, T.D., Petit, R.A., Joseph, S.J., Alam, M.T., Weil, M.R., Ahmad, M., Bhimani, R., Vuong, J.S., Haase, C.P., Webb, D.H., Tan, M., Dove, A.D.M. 2017. Draft sequencing and assembly of the genome of the world’s largest fish, the whale shark: *Rhincodon typus* Smith 1828. BMC Genomics, 18, 532. https://doi.org/10.1186/s12864-017-3926-9

Reid, K., Bell, M.A., Veeramah, K.R., 2021. Threespine Stickleback: A Model System For Evolutionary Genomics. Annu Rev Genomics Hum Genet. 22, 357–383. https://doi.org/10.1146/annurev-genom-111720-081402

Robledo, D., Hermida, M., Rubiolo, J.A., Fernández, C., Blanco, A., Bouza, C., Martínez, P., 2017. Integrating genomic resources of flatfish (Pleuronectiformes) to boost aquaculture production. Comp Biochem Physiol Part D Genomics Proteomics. 21, 41–55. https://doi.org/10.1016/j.cbd.2016.12.001

Sampaio, L.A., Bianchini, A., 2002. Salinity effects on osmoregulation and growth of the euryhaline flounder *Paralichthys orbignyanus*. J. Exp. Mar. Biol. Ecol. 269, 187–196. https://doi.org/10.1016/S0022-0981(01)00395-1

Sampaio, L.A., Freitas, L.S., Okamoto, M.H., Louzada, L.R., Rodrigues, R.V., Robaldo, R.B., 2007. Effects of salinity on Brazilian flounder *Paralichthys orbignyanus* from fertilization to juvenile settlement. Aquaculture 262, 340–346. https://doi.org/10.1016/j.aquaculture.2006.09.046

Sarropoulou, E., Fernandes, J.M, 2011. Comparative genomics in teleost species: Knowledge transfer by linking the genomes of model and non-model fish species. Comp. Biochem. Physiol. Part D 6, 92–102. https://doi.org/10.1016/j.cbd.2010.09.003

Seikai, T., 2002. Flounder Culture and Its Challenges in Asia. Reviews in Fisheries Science, 10, 421–432. https://doi.org/10.1080/20026491051721

Seitz, A.C. Robin, N. Gibson, (Ed.), 2008. Flatfishes: Biology and Exploitation. Rev Fish Biol Fisheries 18, 249–250. https://doi.org/10.1007/s11160-007-9072-8

Schmierder, R., Edwards, R., 2011. Quality control and preprocessing of metagenomic datasets. Bioinformatics, 27, 863–64. https://doi.org/10.1093/bioinformat-ics/btr026

Shao, C., Bao, B., Xie, Z., Chen, X., Li, B., Jia, X., Yao, Q., Ortí, G., 2017. The genome and transcriptome of Japanese flounder provide insights into flatfish asymmetry. Nature Genetics, 49: 119–124. https://doi.org/10.1038/ng.3732

Shao, F., Han, M., Peng, Z., 2019. Evolution and diversity of transposable elements in fish genomes. Sci Rep 9, 15399. https://doi.org/10.1038/s41598-019-51888-1

Slater, G.S., Birney, E., 2005. Automated generation of heuristics for biological sequence comparison. BMC Bioinformatics, 6, 31. https://doi.org/10.1186/1471-2105-6-31

Tarailo-Graovac, M., Chen, N., 2009. Using RepeatMasker to identify repetitive elements in genomic sequences. Curr. Protoc. Bioinformatics 25: 4.10.1– 4.10.14. https://doi.org/10.1002/0471250953.bi0410s25

Volff, J.N., 2005. Genome evolution and biodiversity in teleost fish. Heredity, 94(3), 280–294. https://doi.org/10.1038/sj.hdy.6800635

Waterhouse, R.M., Seppey, M., Simão, F.A., Manni, M., Ioannidis P., Klioutchnikov, G., Kriventseva, E.V., Zdobnov, E.M., 2018. BUSCO Applications from Quality Assessments to Gene Prediction and Phylogenomics. Mol. Biol. Evol. 35, 543–548. https://doi.org/10.1093/molbev/msx319

Xu, X.W., Shao, C.W., Xu, H., Zhou, Q., You, F., Wang, N., Li, W.L., Li, M., Chen, S.L., 2020. Draft genomes of female and male turbot Scophthalmus maximus. Sci Data, 12;7(1): 90. https://doi.org/10.1038/s41597-020-0426-6

Yuan, Z., Liu, S., Zhou, T.; Tian, C.; Bao, L., Dunham, R.; Liu, Z., 2018. Comparative genome analysis of 52 fish species suggests differential associations of repetitive elements with their living aquatic environments. BMC Genomics, 19, 141. https://doi.org/10.1186/s12864-018-4516-1

Zhang, Q., Edwards, S.V., 2012. The evolution of intron size in amniotes: A role for powered flight? Genome Biol. Evol, 4, 1033–1043. https://doi.org/10.1093/gbe/evs070

Zhao, A., Guo, H., Jia, L., Guo, B., Zheng, D., Liu, S., Zhang, B., 2021. Genome assembly and annotation at the chromosomal level of first Pleuronectidae: *Verasper variegatus* provides a basis for phylogenetic study of Pleuronectiformes. Genomics, 113, 717–726. https://doi.org/10.1016/j.ygeno.2021.01.024

Zou, Q., Guo, J., Ju, Y., Wu, M., Zeng, X., Hong., 2015. Improving tRNAscan-SE annotation results via Ensemble classifiers. Mol Inform, 34, 761–770. https://doi.org/10.1002/minf.201500031

