## Supplementary figures and images for "*De novo* assembly of the black flounder genome. Why do pleuronectiformes have such a small genome size?"

### Supplementary Figure S1

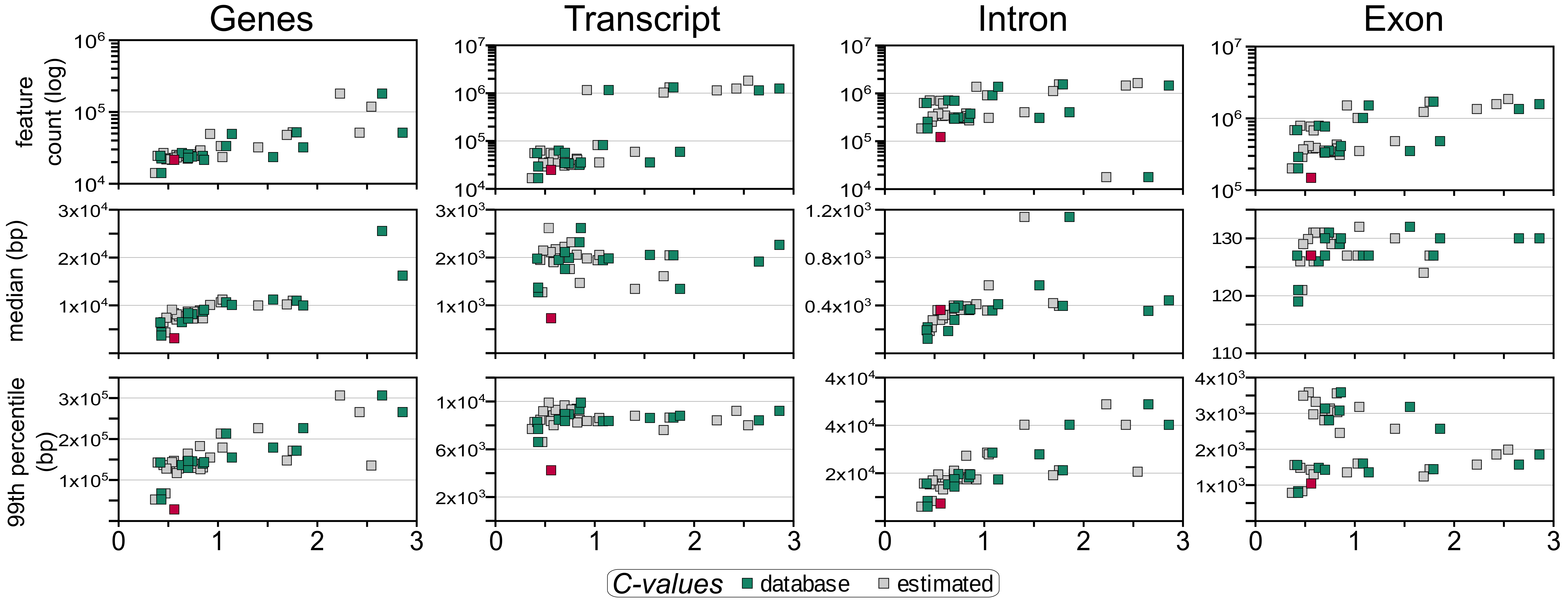

### Supplementary Figure S2

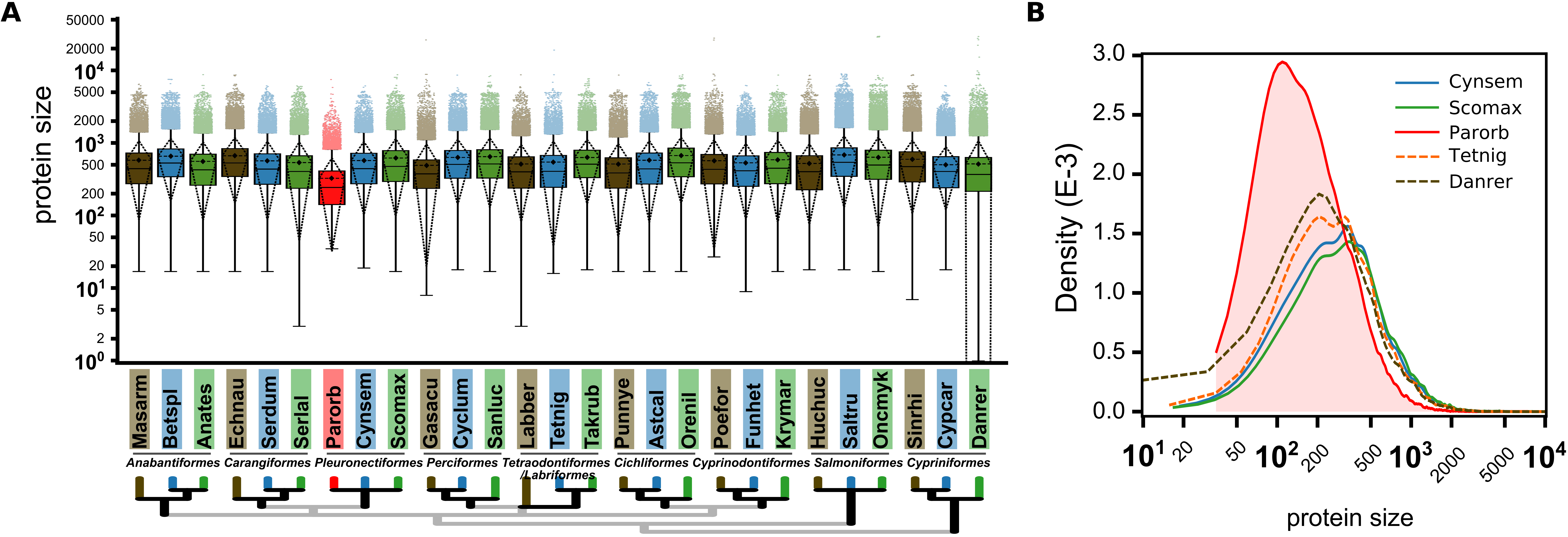

### Supplementary Figure S3

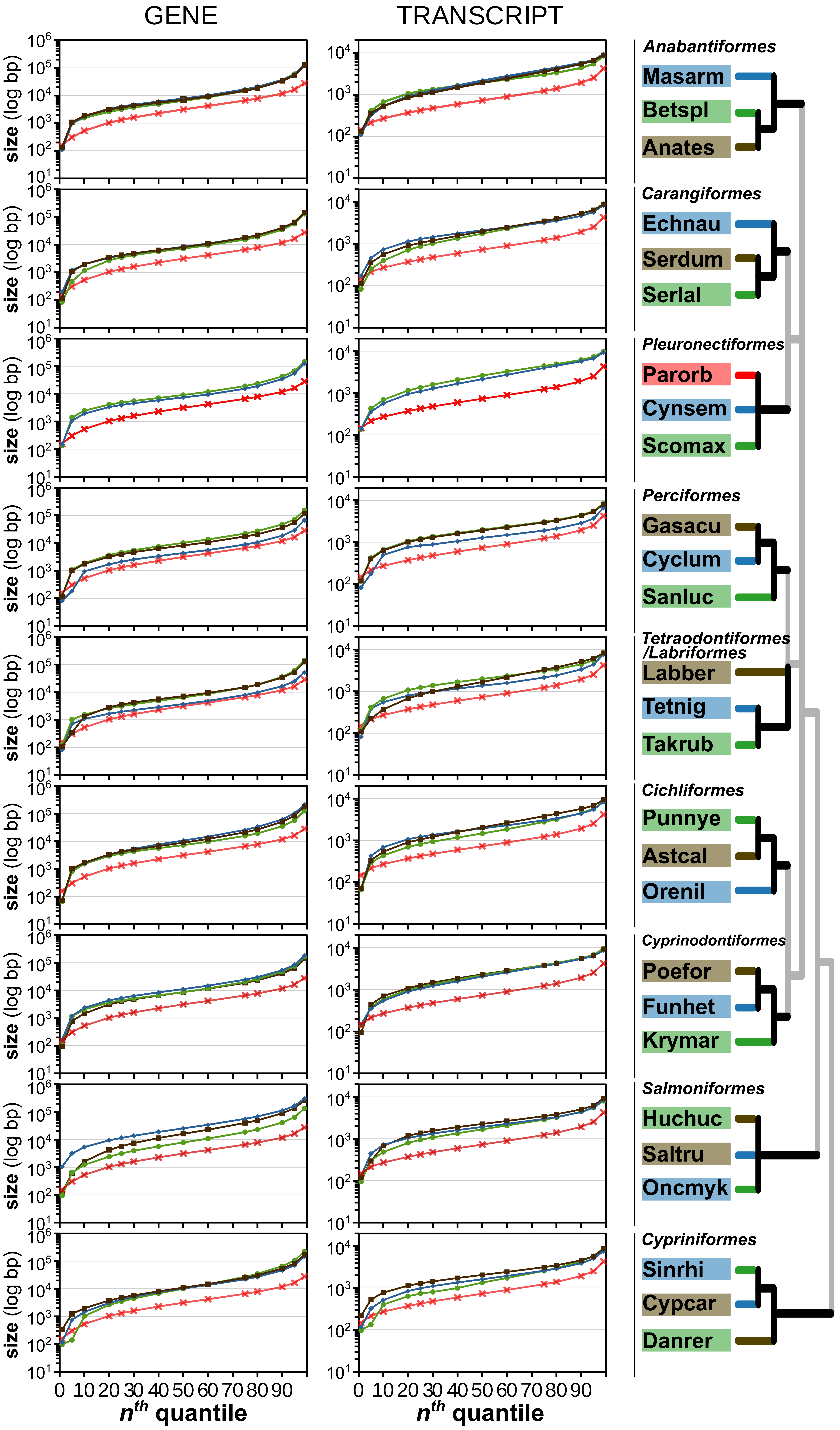

### Supplementary Figure S4

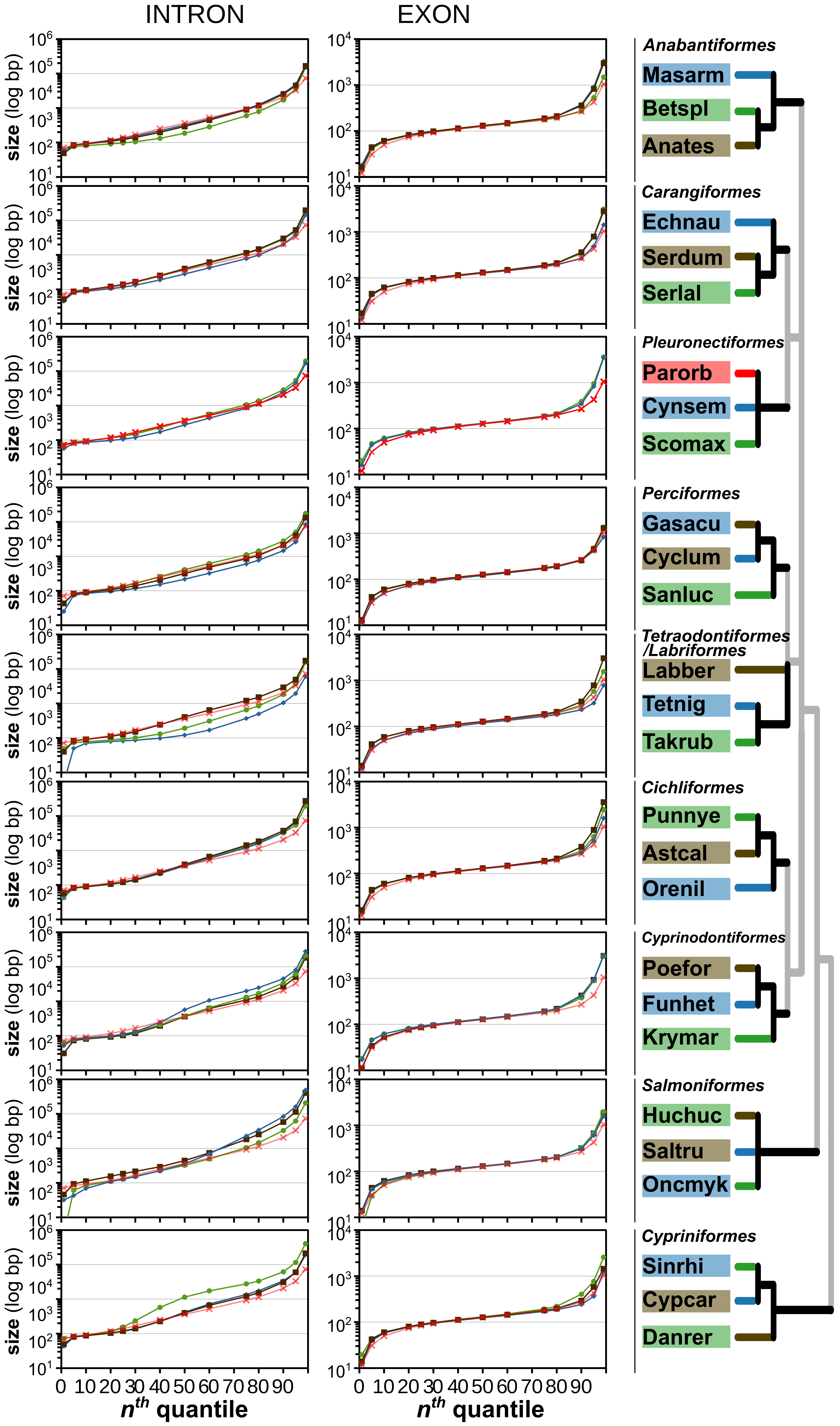
